# Molecular Characterization in 3D Structure of MicroRNA Expressed in Leprosy

**DOI:** 10.1101/2022.02.11.480165

**Authors:** Luís Jesuino de Oliveira Andrade, Humberto Barreto de Jesus, Luís Matos de Oliveira, Moema Farias de Oliveira, Laura Farias Caricchio Santana, Gabriela Correia Matos de Oliveira

## Abstract

**Introduction:** Hansen’s disease, or leprosy, is a major public health problem in developing countries, caused by Mycobacterium leprae, and affecting the skin and peripheral nerves. However, M. leprae can also affect bone tissue, mucous membranes, liver, eyes, and testicles, producing a variety of clinical phenotypes. MicroRNAs (miRNAs) have been expressed in the various clinical forms of leprosy and could potentially be used for its diagnosis.

**Objective:** *In silico* design of the molecular structure of miRNAs expressed in leprosy.

**Method:** We performed a nucleotide sequence search of 17 miRNAs expressed in leprosy, designing *in silico* the molecular structure of the following miRNAs: miRNA-26a, miRNA-27a, miRNA-27b, miRNA-29c, miRNA-34c, miRNA-92a-1, miRNA-99a-2, miRNA-101-1, miRNA-101-2, miRNA-125b-1, miRNA-196b, miRNA-425-5p, miRNA-452, miRNA-455, miRNA-502, miRNA-539, and miRNA-660. We extracted the nucleotides were from the GenBank of National Center for Biotechnology Information genetic sequence database. We aligned the extracted sequences with the RNA Folding Form, and the three-dimensional molecular structure design was performed with the RNAComposer.

**Results:** We demonstrate the nucleotide sequences, and molecular structure projection of miRNAs expressed in leprosy, and produces a tutorial on the molecular model of the 17 miRNAs expressed in leprosy through *in silico* projection processing of their molecular structures.

**Conclusion:** We demonstrate *in silico* design of selected molecular structures of 17 miRNAs expressed in leprosy through computational biology.

## INTRODUCTION

Hansen’s disease, or leprosy, is a major public health problem in developing countries, caused by Mycobacterium leprae (M. leprae), and affecting the skin and peripheral nerves. However, M. leprae can also affect bone tissue, mucous membranes, liver, eyes, and testicles, producing a variety of clinical phenotypes.^1^ About 200,000 new cases were reported in 2017 from almost 150 countries, and migration due to globalization tends to expand the number of cases bringing leprosy to countries where leprosy is not endemic.^2^

Leprosy diagnosis is reasoned on the type and number of lesions, nerve commitment, bacillary load, and histopathology exam. Thus, the bacilloscopy and biopsy of skin from an active lesion, it is considered as “gold standard” in leprosy diagnosis.^3^

The miRNAs has between 19 and 25 nucleotides and are short noncoding RNAs that intervene in post-transcription command of gene manifestation in multicellular beings changing the stableness in system of translation, and consequently miRNAs degradation or inactivation.^4^

MicroRNAs (miRNAs) are relevant elements for host-pathogen interconnection and have been recognized as biological markers of several infectious diseases. Recent information demonstrate that the M. leprae controls the miRNA configuration at the leprosy lesion site interfering with host antimicrobial response.^5^

Studies and understanding about the function and structure of miRNAs have grown exponentially in recent years. In particular, the use of bioinformatics, with careful examination of nucleotide sequences providing essential elements for molecular patterning. This study aims to construct computationally the molecular structure of 17 miRNAs expressed in leprosy, and to produce a tutorial of their modeling.

## METHODS

We performed a nucleotide sequence search of 17 miRNAs expressed in leprosy, performing *in silico* molecular structure design of the most expressed miRNAs in leprosy lesions based on literature review studies: miRNA-26a, miRNA-27a, miRNA-27b, miRNA-29c, miRNA-34c, miRNA-92a-1, miRNA-99a-2, miRNA-101-1, miRNA-101-2, miRNA-125b, miRNA-196b, miRNA-425, miRNA-452, miRNA-455, miRNA-502, miRNA-539, and miRNA-660.

We rescued the nucleotides from the GenBank of National Center for Biotechnology Information genetic sequence database (https://www.ncbi.nlm.nih.gov/). We aligned the extracted sequences with the RNA Folding Form (http://www.unafold.org/), and the three-dimensional molecular structure design was performed with the RNAComposer (https://rnacomposer.cs.put.poznan.pl/), as employed to design the molecular configurations. We proceeded with the development of a tutorial on the molecular model of the 17 miRNAs described above by performing *in silico* their molecular arrangement.

### Nucleotide search and sequence analysis

GenBank is public domain nucleotide sequence assessment software accessible through the electronic address: https://www.ncbi.nlm.nih.gov/genbank/submit/. The GenBank disposes of numerous nucleotide research algorithms and a great diversity of nucleotide algorithms can be used to investigate the different databases of distinct sequences. Furthermore, GenBank possesses the Nucleotide, Genome Survey Sequence (GSS) and the Expressed Sequence Tag (EST) including nucleic acid sequences. The references contained are derived from the distribution of the number of sequences from GenBank

### Molecular model construction

The function and constitution of amino acids and proteins are composed of nucleotide sequences and structure forecast until then is a significant problem, with a vast search for high-resolution structure methods.

Modeling by homology is at present the most precise method computational method to produce trustworthy structural models and is generally utilized in numerous biological uses.

### Designing the structure of RNA

The RNAComposer system presents an interactive proposal for the fully automated prediction of RNA three-dimensional (3-D) structures. The system is based on the precept of automatic translation and works on the RNA FRABASE database as the dictionary concerning the secondary structure of the RNA and the components of the tertiary structure.

RNAComposer works interactively, allowing you to work with an RNA molecule of up to 500 nucleotide residues, resulting in only one model of the RNA three-dimensional structure. Batch mode, on the other hand, is designed for automated modeling of RNA structures of up to 500 nucleotide residues, and up to 10 RNAs sequences can be used. The RNAComposer package is free of charge available for public use and download, from homepage at: https://rnacomposer.cs.put.poznan.pl/.

## RESULTS

We showed a search for nucleotide sequence and the drawing of molecular structure of following miRNA expressed in lesions due to leprosy: miR-26a, miR-27a, miR-27b, miR-29c, miR-34c, miR-92a-1, miR-99a-2, miR-101-1, miR-101-2, miR-125b-1, miR-196b, miR-425, miR-452, miR-455-3p, miR-502-3p, miR-539, and miR-660. We produced a tutorial on molecular model of the 17 miRNAs expressed in lesions due to leprosy through of processing *in silico* projection of their molecular structures.

### Nucleotide sequence of miRNA-26a

To construct the structure of miRNA-26a-1 the nucleotide sequences with the NCBI identifier code: AJ421747. were used under the FASTA format obtained from GenBank. The Homo sapiens microRNA miR-26a was planned to encode a 22 bp linear ncRNA. Homo sapiens microRNA 26a (MIR26A), microRNA analysis is demonstrated in Figure 1.

**Figure 1.**
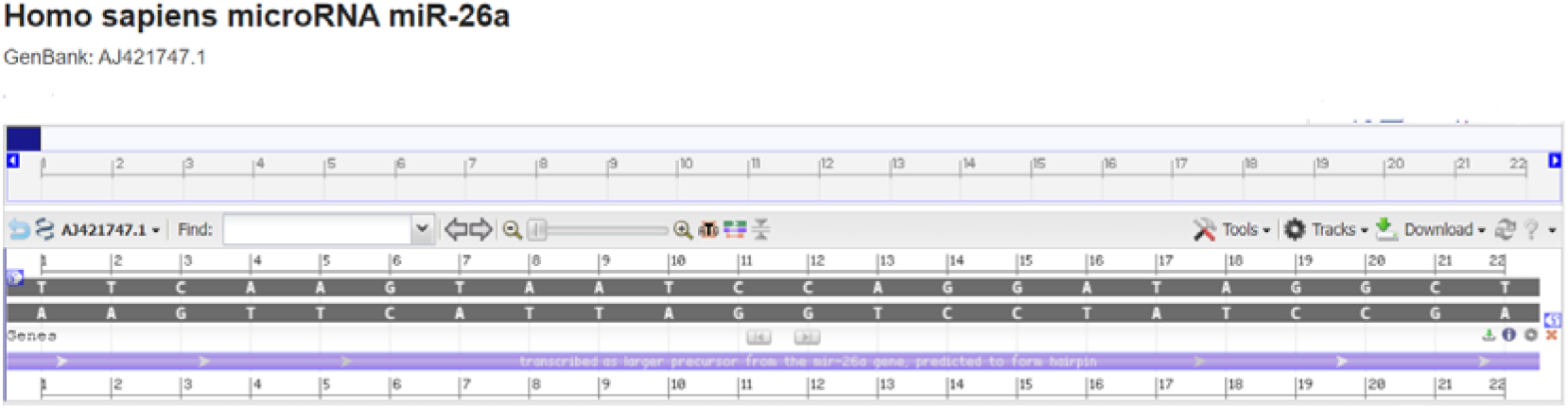
Homo sapiens microRNA miR-26a - *model-template aligment*. **Source**: https://www.ncbi.nlm.nih.gov/nuccore/AJ421747.1?report=graph

### Molecular model of miRNA-26a

Nucleotide sequences of Homo sapiens microRNA 26a (MIR26A) were acquired employing FASTA format sequence: UUCAAGUAAUCCAGGAUAGGCU and secondary structure: ….(((.(((…)))..)))); modeling was performed employing the RNAComposer, optimized and adjusted for alignment between structural templates and miRNA-26a nucleotide. Based on sequence alignment between the template structure and miRNA-26a nucleotide, a structural model was built for the nucleotide in question. So, employing RNAComposer of comparative nucleotide modeling we generated a homology model of microRNA 26a (MIR26A), demonstrated in Figure 2 with CPK spacefill style structure in pdb format.

**Figure 2.**
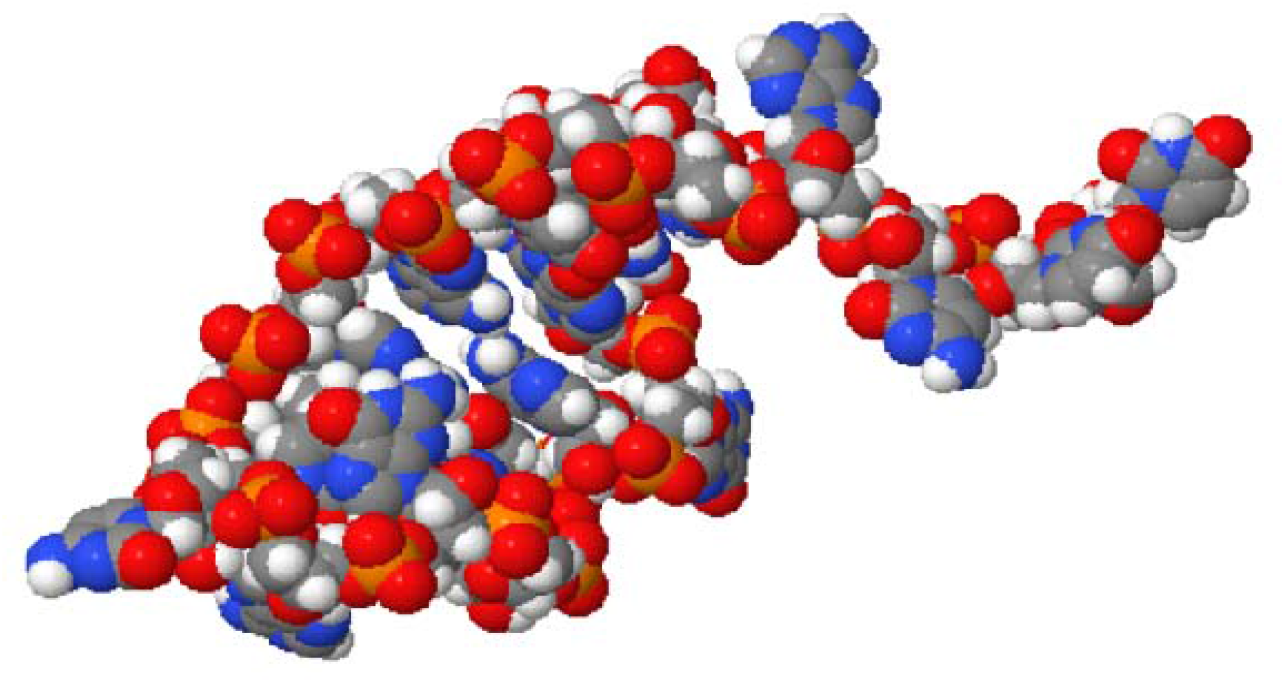
Homology model of the Homo sapiens microRNA26a (MIR26A). **Source**: https://rnacomposer.cs.put.poznan.pl/;jsessionid=2F086BF953870CD041DC6D65AAFBAABE

### Nucleotide sequence of miRNA-27a

The reconstruction of miRNA-27a was performed from a nucleotide sequence archive in FASTA format obtained in GenBank database with the identifier code NCBI Reference Sequence: NR_029501.1. The miRNA-27a was planned to encode a 78 bp linear ncRNA. All coded sequences selected in FASTA format, used the annotation of the NCBI - Graphics. The Homo sapiens microRNA 27a (MIR27A), analysis is shown in Figure 3.

**Figure 3.**
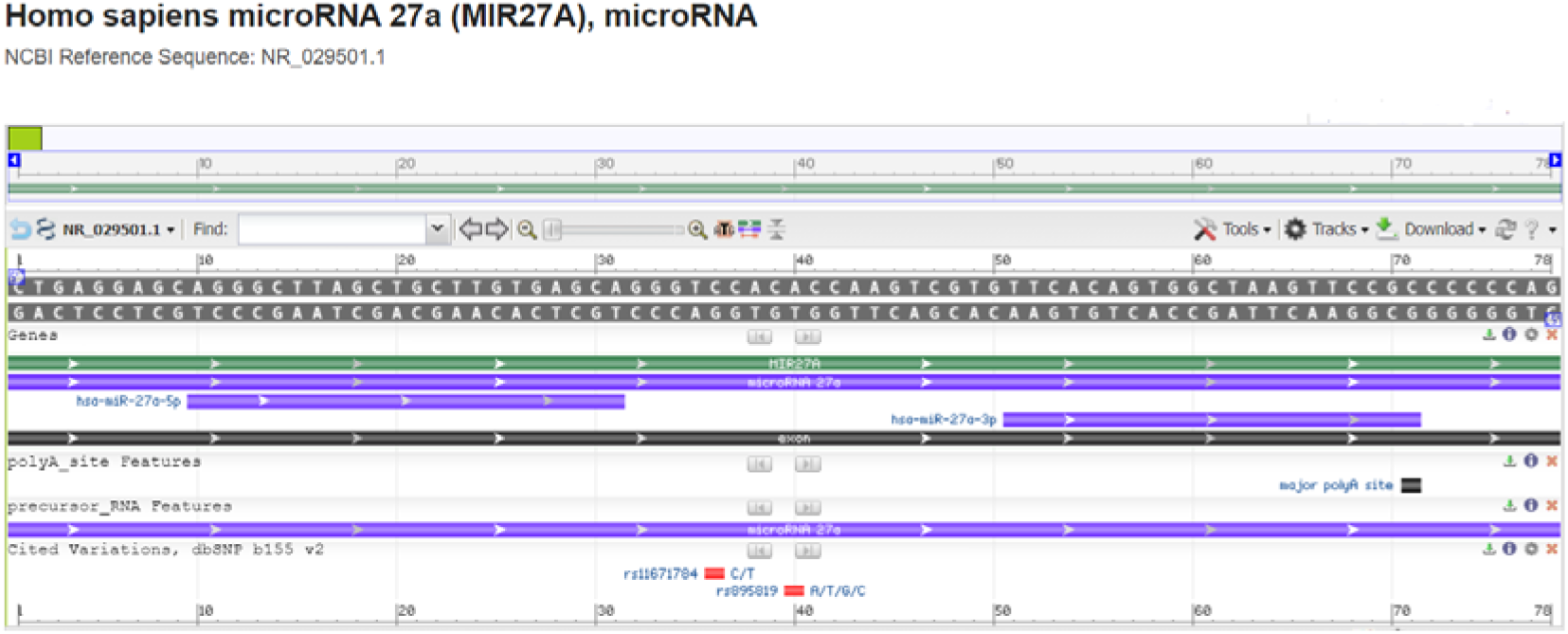
Homo sapiens microRNA 27a (MIR27A), microRNA - *model-template aligment*. **Source**: https://www.ncbi.nlm.nih.gov/nuccore/NR_029501.1?report=graph

### Molecular model of microRNA 27a

Based on the sequence alignment between the Homo sapiens microRNA 27a nucleotide sequence: CUGAGGAGCAGGGCUUAGCUGCUUGUGAGCAGGGUCCACACCAAGUCGUGUUCACAGUGGCUAAGUUCCGCCCCCCAG and secondary structure: (((.((.((.((((((((((((.((((((((.(((….)))……)))))))))))))))))))).)).)).))) and the template structure, the structural template for microRNA 27a was produced. Assessment tools were used para measure the reliability of the designed structure. Thereby, using the RNAComposer comparative nucleotide modeling server, we generate a homology model of the Homo sapiens microRNA 27a, with Cartoon-style structure in pdb format (Figure 4).

**Figure 4.**
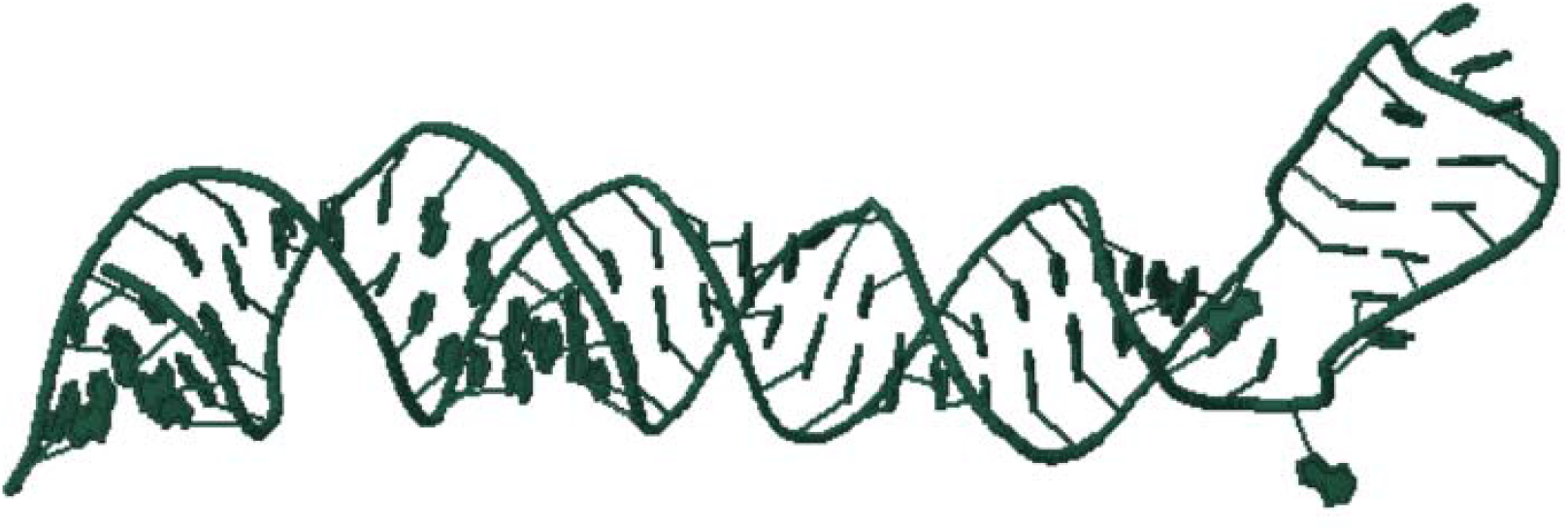
Homology model of the Homo sapiens microRNA 27a (MIR27A). **Source**: https://rnacomposer.cs.put.poznan.pl/

### Nucleotide sequence of miRNA-27b

To construct the structure of Homo sapiens microRNA 27b, the nucleotide sequences with the NCBI identifier code: NR_029665.1 were used under the FASTA format obtained from GenBank. The miRNA-27b was planned to encode a 97 bp linear ncRNA. Homo sapiens microRNA 27b (MIR27B), microRNA analysis is demonstrated in Figure 5.

**Figure 5.**
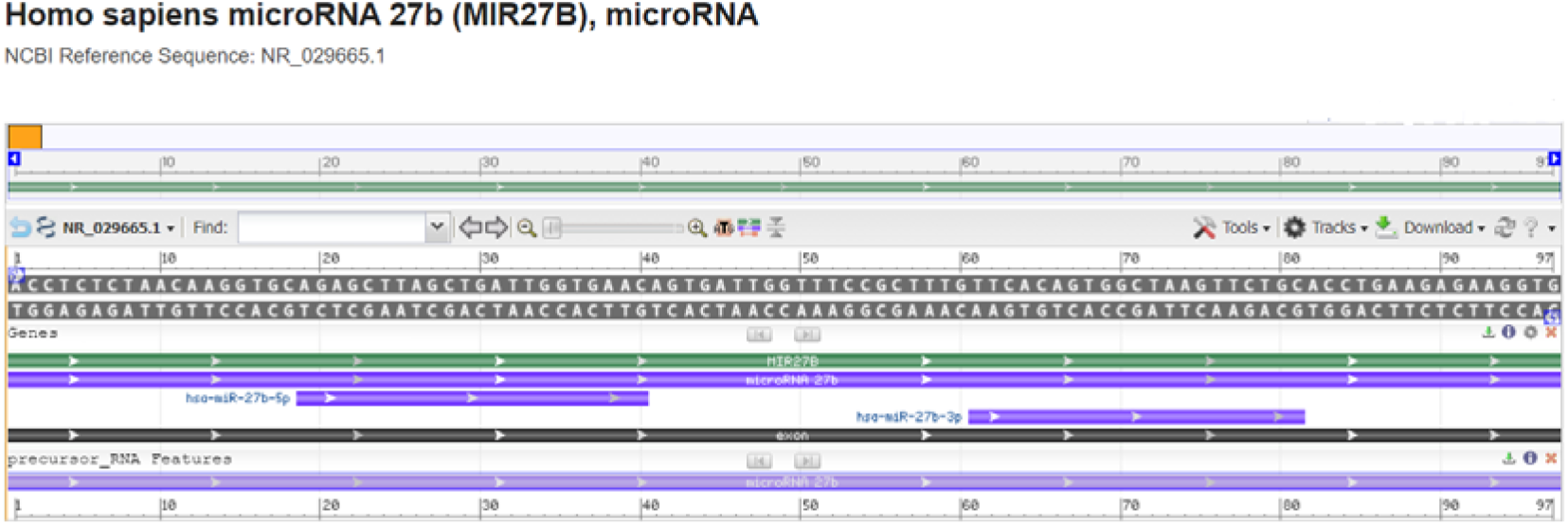
Homo sapiens microRNA 27b (MIR27B), microRNA - *model-template aligment*. **Source**: https://www.ncbi.nlm.nih.gov/nuccore/NR_029665.1?report=graph

### Molecular model of microRNA 27b

Nucleotide sequences of Homo sapiens microRNA 27b (MIR27B) were acquired employing FASTA format sequence:ACCUCUCUAACAAGGUGCAGAGCUUAGCUGAUUGGUGAACAGUGAUUGGUUUCCGCUUUGUUCACAGUGGCUAAGUUCUGCACCUGAAGAGAAGGU and secondary structure: ((((((((….((((((((((((((((((….((((((((….(((…)))..))))))))..))))))))))))))))))..))))).))).; modeling was performed employing the RNAComposer, optimized and adjusted for alignment between structural templates and microRNA 27b nucleotide. Based on sequence alignment between the template structure and microRNA nucleotide, a structural model was built for the nucleotide in question. So, employing RNAComposer of comparative nucleotide modeling we generated a homology model of Homo sapiens microRNA 27b (MIR27B), demonstrated in Figure 6 with CPK spacefill style structure in pdb format.

**Figure 6.**
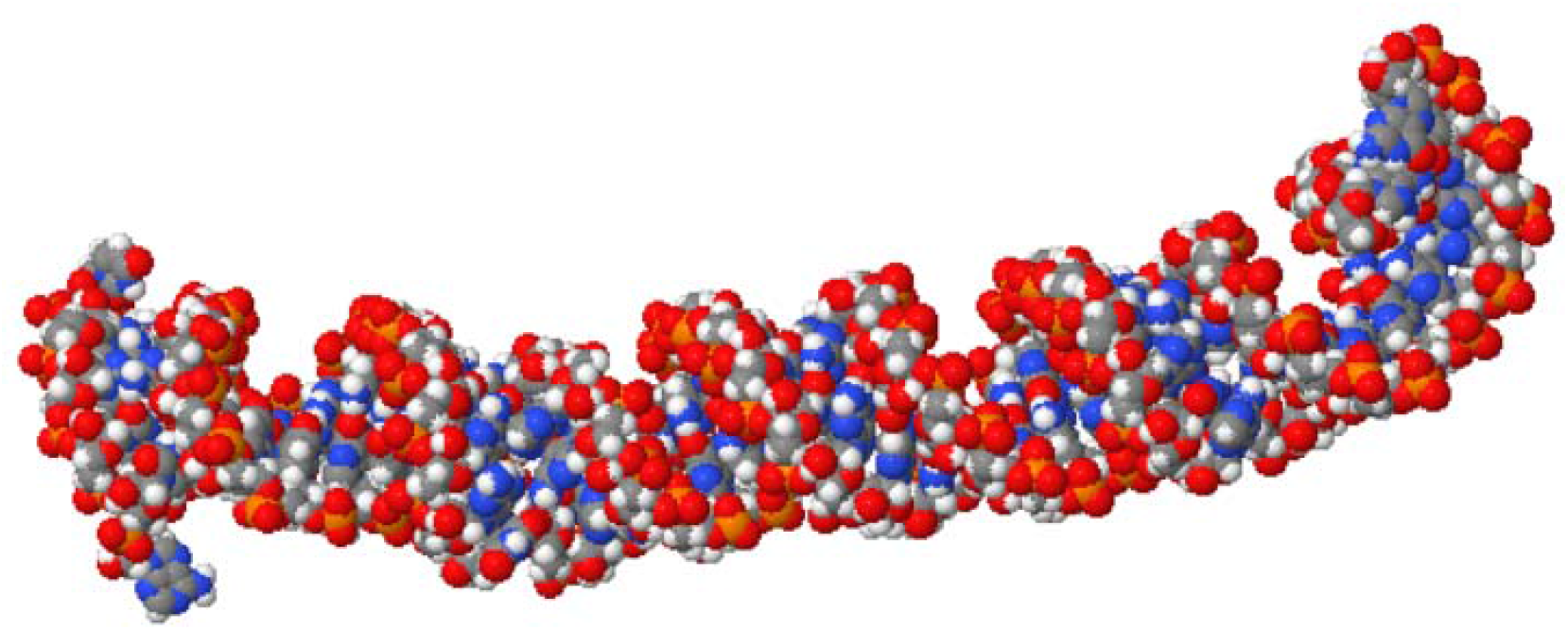
Homology model of the Homo sapiens microRNA 27b (MIR27B) **Source**: https://rnacomposer.cs.put.poznan.pl/

### Nucleotide sequence of miRNA-29c

The reconstruction of miRNA-29c was performed from a nucleotide sequence archive in FASTA format obtained in GenBank database with the identifier code NCBI Reference Sequence: NR_029832.1. The miRNA-29c was planned to encode a 88 bp linear ncRNA. All coded sequences selected in FASTA format, used the annotation of the NCBI - Graphics - Homo sapiens microRNA 29c (MIR29C). The microRNA-29c analysis is shown in Figure 7.

**Figure 7.**
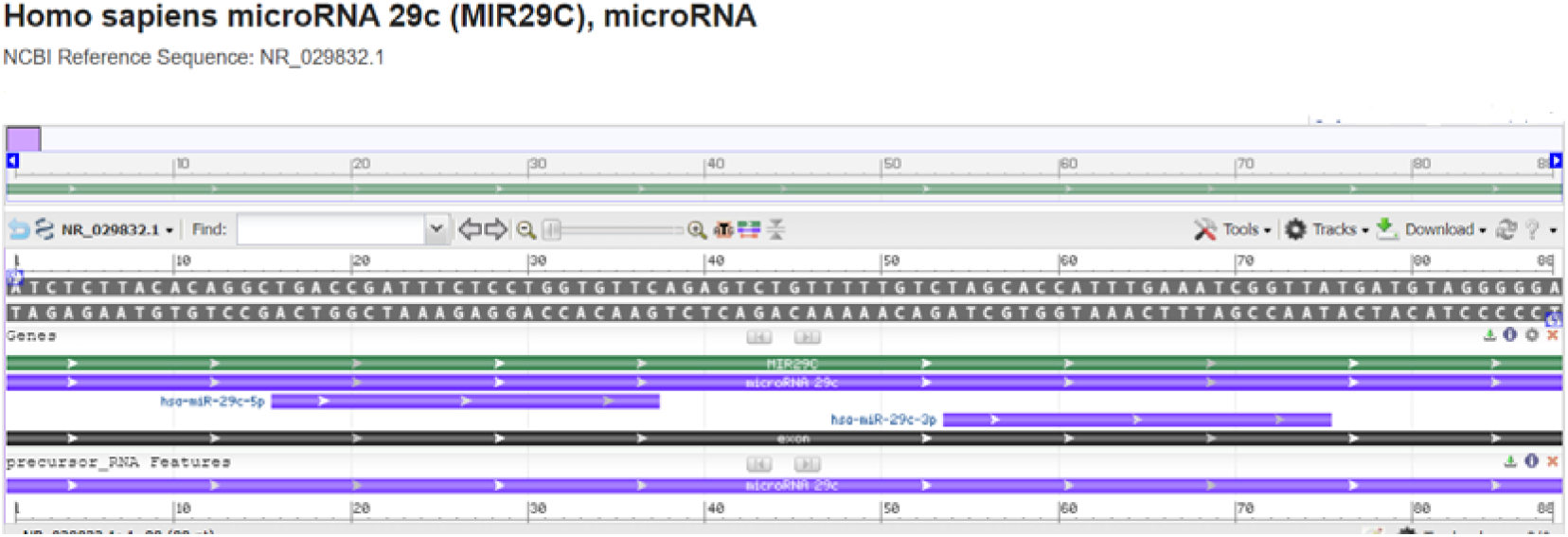
Homo sapiens microRNA 29c (MIR29C), microRNA - *model-template aligment*. **Source**: https://www.ncbi.nlm.nih.gov/nuccore/NR_029832.1?report=graph

### Molecular model of microRNA-29c

Based on the sequence alignment between the Homo sapiens microRNA 29c nucleotide sequence: AUCUCUUACACAGGCUGACCGAUUUCUCCUGGUGUUCAGAGUCUGUUUUUGUCUAGCACCAUUUGAAAUCGGUUAUGAUGUAGGGGGA and secondary sequence: .(((((((((((…(((((((((((…(((((((.(((………..))))))))))…))))))))))))).))))))))). and the template structure, the structural template for microRNA-29c was produced. Assessment tools were used para measure the reliability of the designed structure. Thereby, using the RNAComposer comparative nucleotide modeling server, we generate a homology model of the Homo sapiens microRNA 29c with Cartoon-style structure in pdb format (Figure 8).

**Figure 8.**
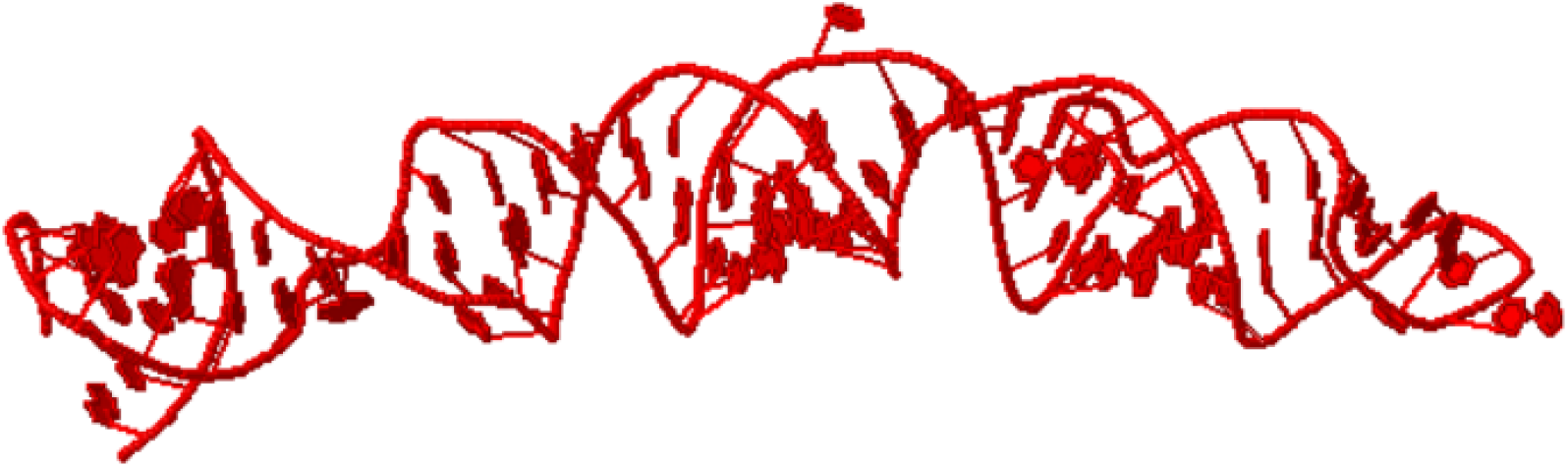
Homology model of the Homo sapiens microRNA 29c (MIR29C), microRNA **Source**: https://rnacomposer.cs.put.poznan.pl/

### Nucleotide sequence of miRNA-34c

To construct the structure of miRNA-34c the nucleotide sequences with the NCBI identifier code: NR_029840.1 were used under the FASTA format obtained from GenBank. The miRNA-21 was planned to encode a 77 bp linear ncRNA. Homo sapiens microRNA 34c (MIR34c) analysis is demonstrated in Figure 9.

**Figure 9.**
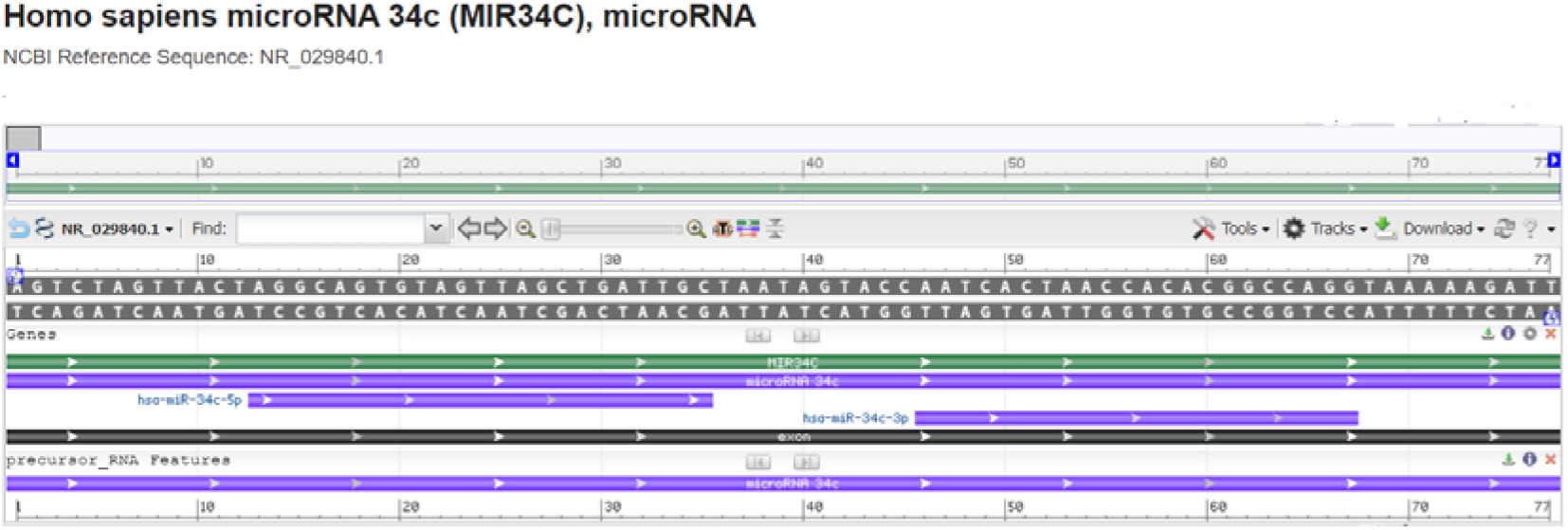
Homo sapiens microRNA 34c (MIR34c) - *model-template aligment*. **Source**: https://www.ncbi.nlm.nih.gov/nuccore/NR_029840.1?report=graph

### Molecular model of miRNA-34c

Nucleotide sequences of Homo sapiens microRNA 34c (MIR34c) were acquired employing FASTA format sequence: AGUCUAGUUACUAGGCAGUGUAGUUAGCUGAUUGCUAAUAGUACCAAUCACUAACCACACGGCCAGGUAAAAAGAUU, and secondary structure: .((((..(((((.(((.((((.(((((.((((((.((….)).))))))))))).)))).))).)))))..)))).; modeling was performed employing the RNAComposer, optimized and adjusted for alignment between structural templates and miRNA-34c nucleotide. Based on sequence alignment between the template structure and miRNA-34c nucleotide, a structural model was built for the nucleotide in question. So, employing RNAComposer of comparative nucleotide modeling we generated a homology model of microRNA 34c (MIR34c), demonstrated in Figure 10 with CPK spacefill style structure in pdb format.

**Figure 10.**
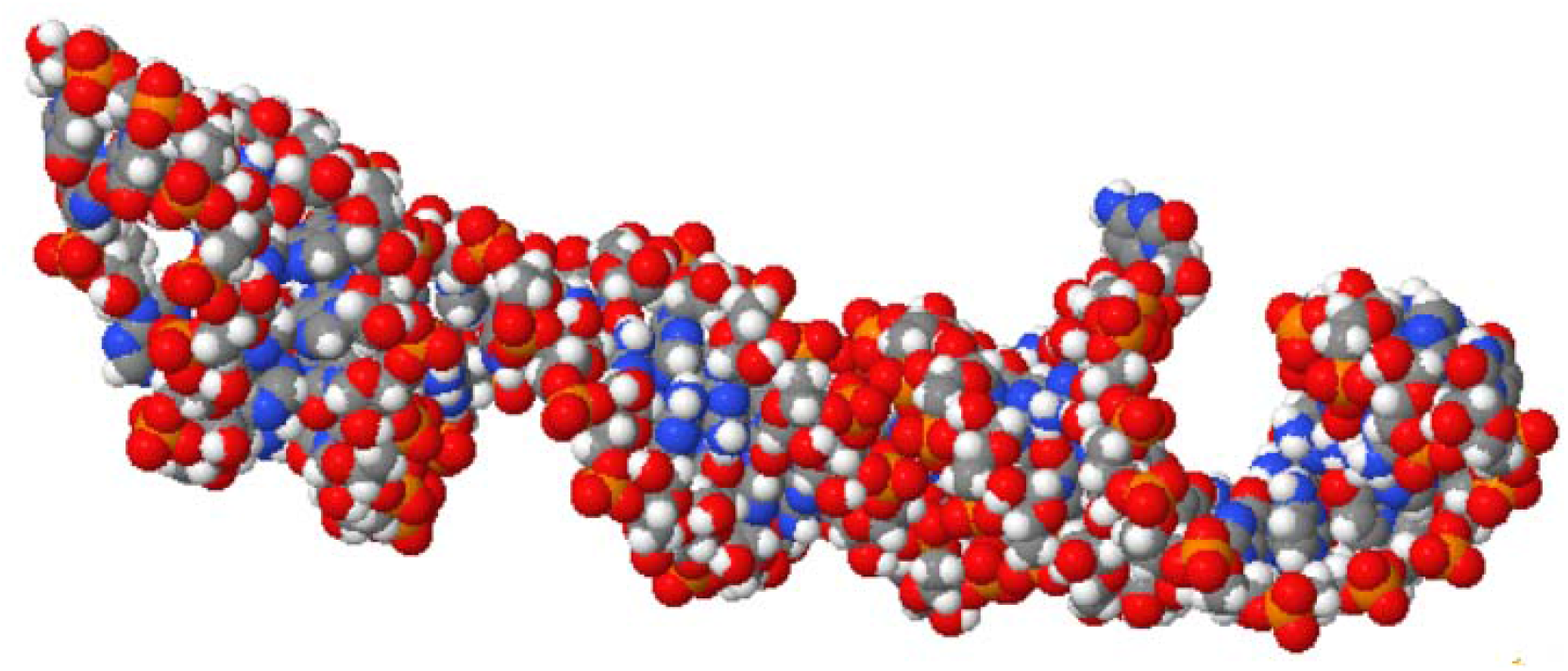
Homology model of the Homo sapiens microRNA 34c (MIR34C), microRNA **Source**: https://rnacomposer.cs.put.poznan.pl/

### Nucleotide sequence of miRNA-92a

The reconstruction of miRNA-92a-1 was performed from a nucleotide sequence archive in FASTA format obtained in GenBank database with the identifier code NCBI Reference Sequence: NR_029508.1. The miRNA-92a-1 was planned to encode a 78 bp linear ncRNA. All coded sequences selected in FASTA format, used the annotation of the NCBI - Graphics. The Homo sapiens microRNA 92a-1 (MIR92A-1), microRNA analysis is shown in Figure 11.

**Figure 11.**
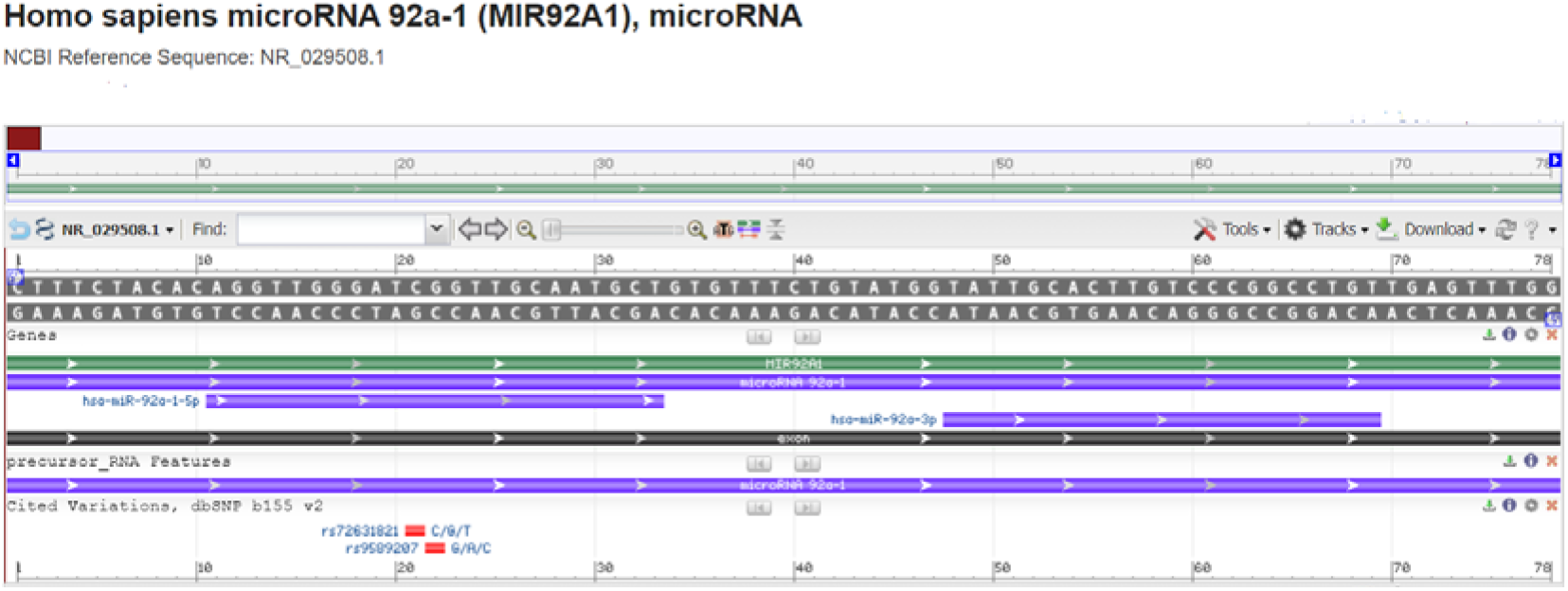
Homo sapiens microRNA 92a-1 (MIR92A-1) - *model-template aligment*. **Source**: https://www.ncbi.nlm.nih.gov/nuccore/NR_029508.1?report=graph

### Molecular model of microRNA-92a-1

Based on the sequence alignment between the Homo sapiens microRNA 92a-1 nucleotide sequence: CUUUCUACACAGGUUGGGAUCGGUUGCAAUGCUGUGUUUCUGUAUGGUAUUGCACUUGUCCCGGCCUGUUGAGUUUGG, and secondary structure: ..(((…((((((((((((.(((.(((((((((((……)))))))))))))).)))))))))))).)))….., and the template structure, the structural template for microRNA 203a was produced. Assessment tools were used para measure the reliability of the designed structure. Thereby, using the RNAComposer comparative nucleotide modeling server, we generate a homology model of the Homo sapiens microRNA 92a-1 (MIR92A1) with Cartoon-style structure in pdb format (Figure 12).

**Figure 12.**
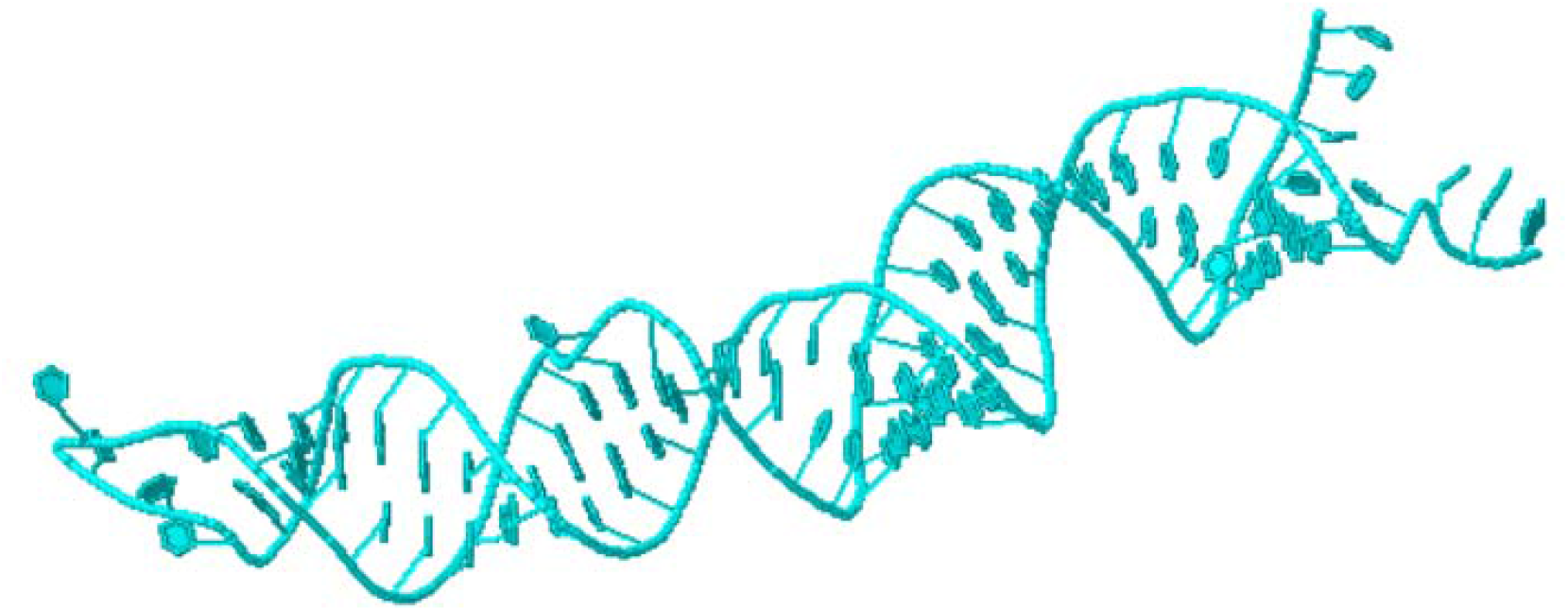
Homology model of the Homo sapiens microRNA 92a-1 (MIR92A1), microRNA **Source**: https://rnacomposer.cs.put.poznan.pl/

### Nucleotide sequence of miRNA-92a-2

To construct the structure of miRNA-92a-2 the nucleotide sequences with the NCBI identifier code: NR_029509.1 were used under the FASTA format obtained from GenBank. The miRNA-92a-2 was planned to encode a 75 bp linear ncRNA. Homo sapiens microRNA 92a-2 (MIR92A2), microRNA, microRNA analysis is demonstrated in Figure 13.

**Figure 13.**
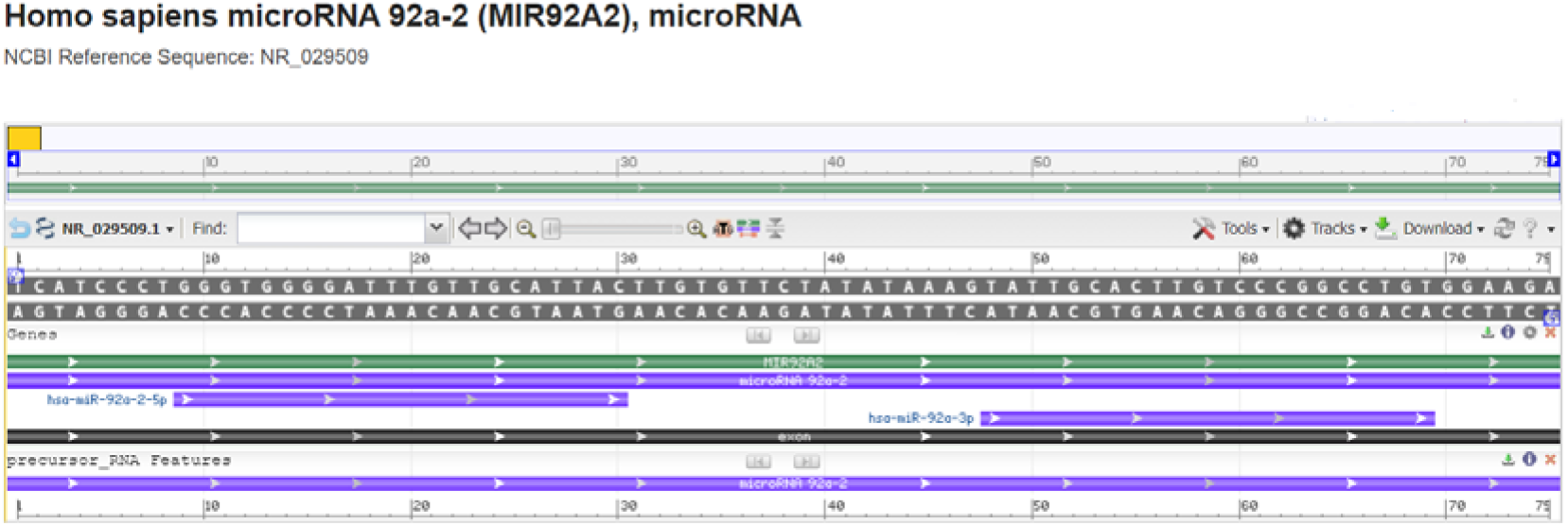
Homo sapiens microRNA 92a-2 (MIR92A2), microRNA - *model-template aligment*. **Source**: https://www.ncbi.nlm.nih.gov/nuccore/NR_029509.1?report=graph

### Molecular model of microRNA-92a-2

Nucleotide sequences of Homo sapiens microRNA 92a-2 (MIR92A2), microRNA were acquired employing FASTA format sequence: UCAUCCCUGGGUGGGGAUUUGUUGCAUUACUUGUGUUCUAUAUAAAGUAUUGCACUUGUCCCGGCCUGUGGAAGA, and secondary structure: ((.(((.(((((.(((((..((.(((.(((((…………))))).)))))..))))).))))).))).)); modeling was performed employing the RNAComposer, optimized and adjusted for alignment between structural templates and miRNA-92a-2 nucleotide. Based on sequence alignment between the template structure and miRNA-92a-2 nucleotide, a structural model was built for the nucleotide in question. So, employing RNAComposer of comparative nucleotide modeling we generated a homology model of microRNA 92a-2 (MIR92A2), with CPK spacefill style structure in pdb format demonstrated in Figure 14.

**Figure 14.**
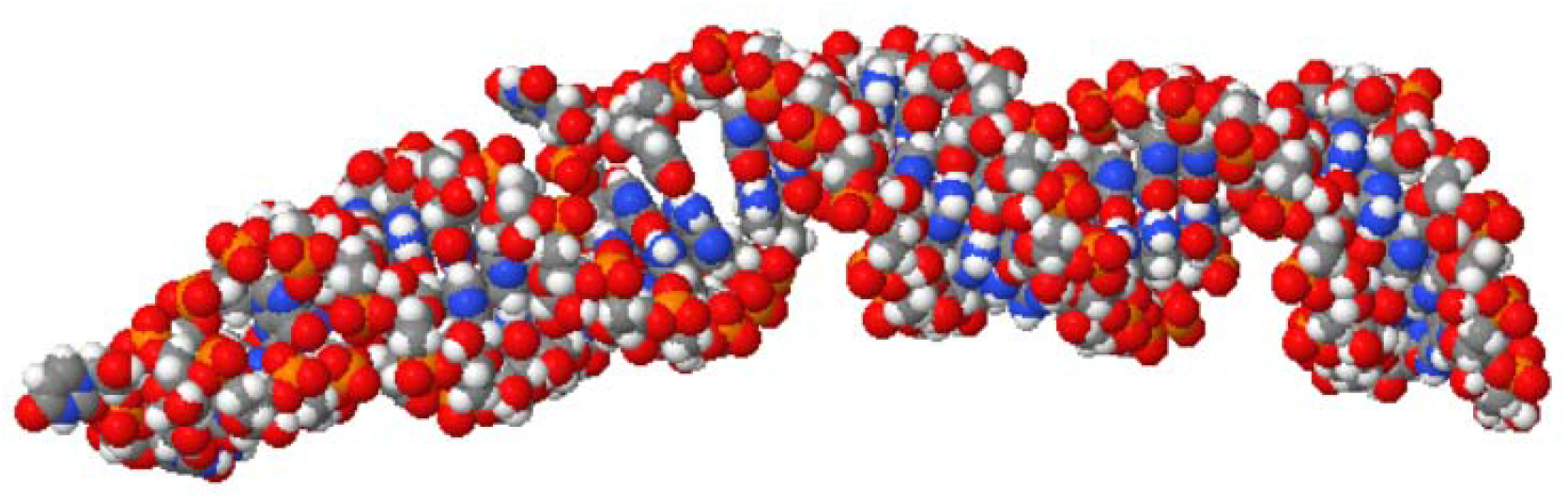
Homology model of the Homo sapiens microRNA 92a-2 (MIR92A2), microRNA **Source**: https://rnacomposer.cs.put.poznan.pl/

### Nucleotide sequence of miRNA-101-1

The reconstruction of miRNA-101-1 was performed from a nucleotide sequence archive in FASTA format obtained in GenBank database with the identifier code NCBI Reference Sequence: NR_029516.1. The miRNA-101-1 was planned to encode a 75 bp linear ncRNA. All coded sequences selected in FASTA format, used the annotation of the NCBI - Graphics. The Homo sapiens microRNA 101-1 (MIR101-1), microRNA analysis is shown in Figure 15.

**Figure 15.**
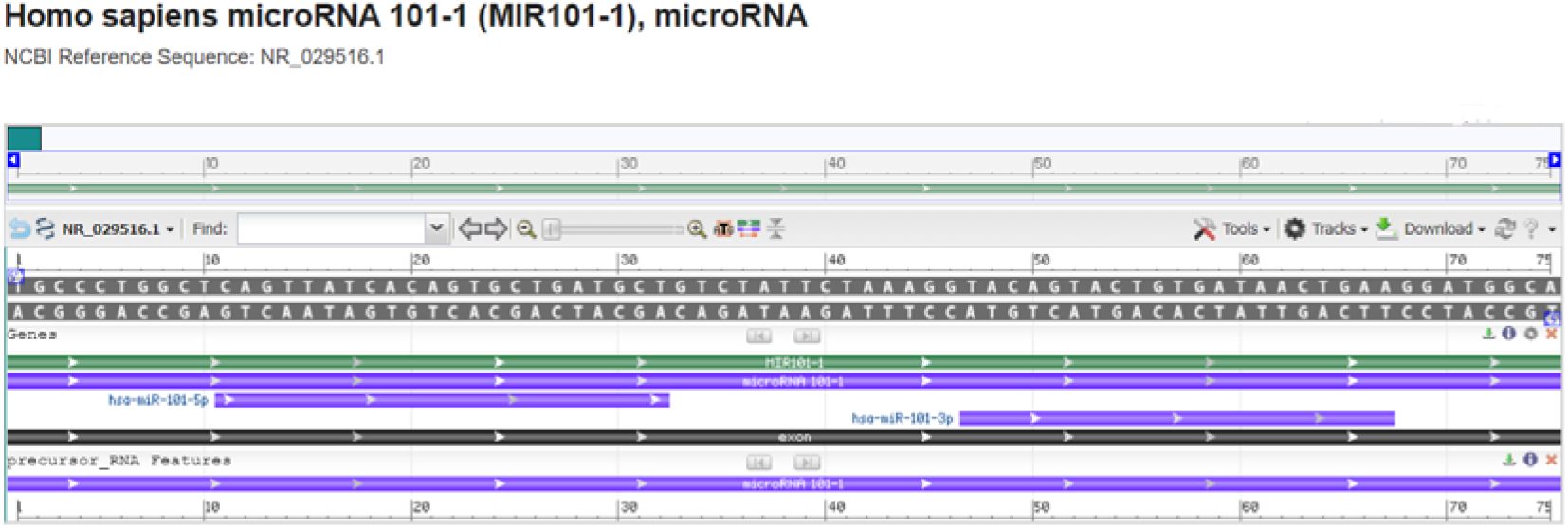
Homo sapiens microRNA 101-1 (MIR101-1), microRNA - *model-template aligment*. **Source**: https://www.ncbi.nlm.nih.gov/nuccore/NR_029516.1?report=graph

### Molecular model of microRNA-101-1

Based on the sequence alignment between the Homo sapiens microRNA-101-1 nucleotide sequence: UGCCCUGGCUCAGUUAUCACAGUGCUGAUGCUGUCUAUUCUAAAGGUACAGUACUGUGAUAACUGAAGGAUGGCA, and secondary structure: ((((((…((((((((((((((((((.((((…………)))))))))))))))))))))).))..)))) and the template structure, the structural template for microRNA-101-1was produced. Assessment tools were used para measure the reliability of the designed structure. Thereby, using the RNAComposer comparative nucleotide modeling server, we generate a homology model of the Homo sapiens microRNA 101-1 (MIR101-1), microRNA with Cartoon-style structure in pdb format (Figure 16).

**Figure 16.**
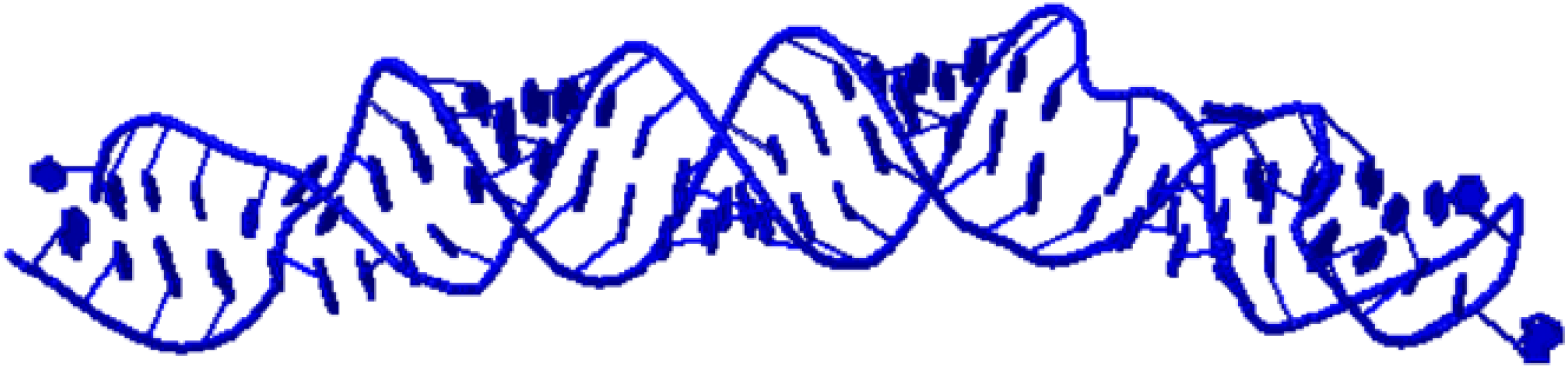
Homology model of the Homo sapiens microRNA 101-1 (MIR101-1), microRNA **Source**: https://rnacomposer.cs.put.poznan.pl/

### Nucleotide sequence of miRNA-101-2

To construct the structure of miRNA-101-2 the nucleotide sequences with the NCBI identifier code: NR_029836.1 were used under the FASTA format obtained from GenBank. The miRNA-101-2 was planned to encode a 79 bp linear ncRNA. Homo sapiens microRNA 101-2 (MIR101-2), microRNA analysis is presented at Figure 17.

**Figure 17.**
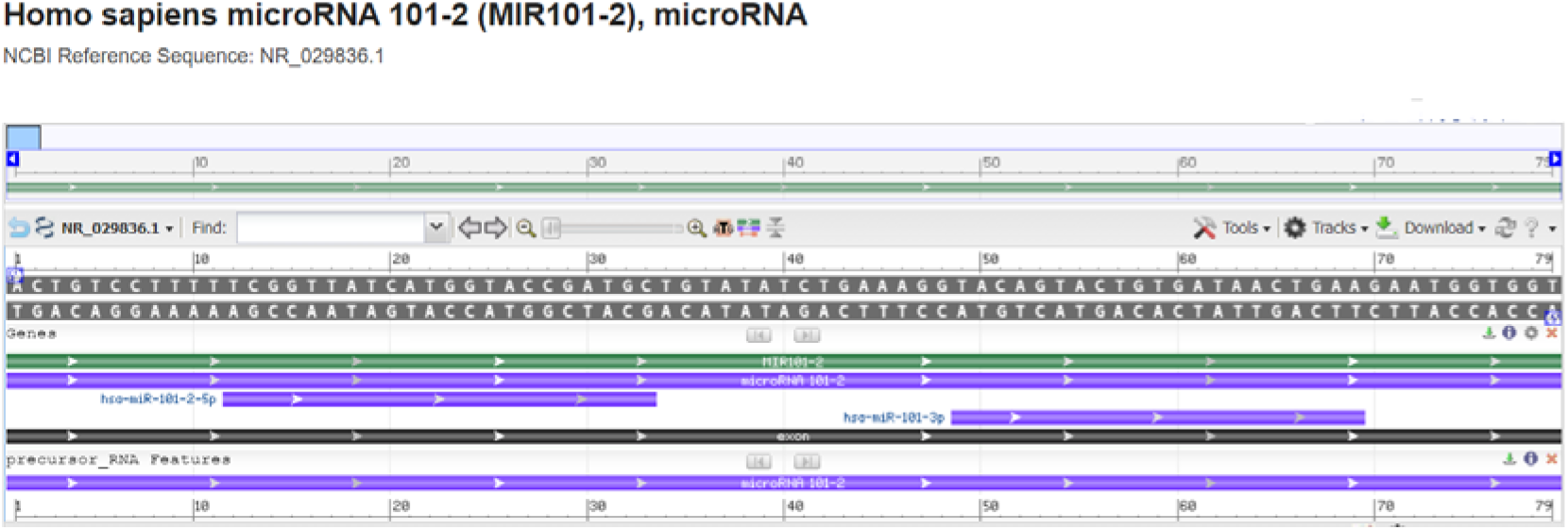
Homo sapiens microRNA 101-2 (MIR101-2), microRNA - *model-template aligment*. **Source**: https://www.ncbi.nlm.nih.gov/nuccore/NR_029836.1?report=graph

### Molecular model of miRNA-101-2

Nucleotide sequences of Homo sapiens microRNA 101-2 (MIR101-2) were acquired employing FASTA format sequence: ACUGUCCUUUUUCGGUUAUCAUGGUACCGAUGCUGUAUAUCUGAAAGGUACAGUACUGUGAUAACUGAAGAAUGGUGGU, and secondary structure: (((.((.((((((((((((((((((……(((((((………))))))))))))))))))))))))).)).))); modeling was performed employing the RNAComposer, optimized and adjusted for alignment between structural templates and miRNA-101-2 nucleotide. Based on sequence alignment between the template structure and miRNA-101-2 nucleotide, a structural model was built for the nucleotide in question. So, employing RNAComposer of comparative nucleotide modeling we generated a homology model of microRNA 101-2 (MIR101-2), with CPK spacefill style structure in pdb format demonstrated in Figure 18.

**Figure 18.**
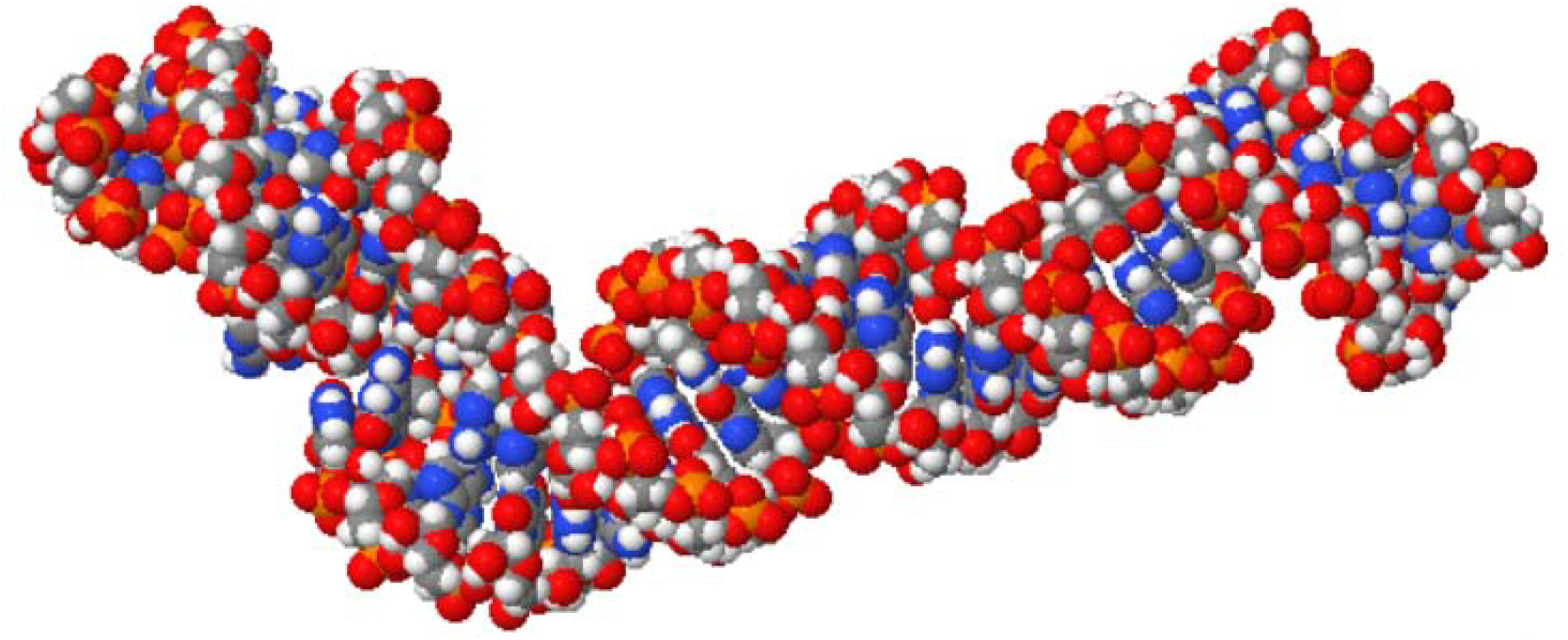
Homology model of the Homo sapiens microRNA 101-2 (MIR101-2). **Source**: https://rnacomposer.cs.put.poznan.pl/;jsessionid=084F1BA99C3734CDB5CB4785BDD7E4F8

### Nucleotide sequence of miRNA-125b-1

The reconstruction of miRNA-125b-1 was performed from a nucleotide sequence archive in FASTA format obtained in GenBank database with the identifier code NCBI Reference Sequence: NR_029671.1. The miRNA-125b-1 was planned to encode a 88 bp linear ncRNA. All coded sequences selected in FASTA format, used the annotation of the NCBI - Graphics. The Homo sapiens microRNA 125b-1 (MIR125B1), microRNA analysis is shown in Figure 19.

**Figure 19.**
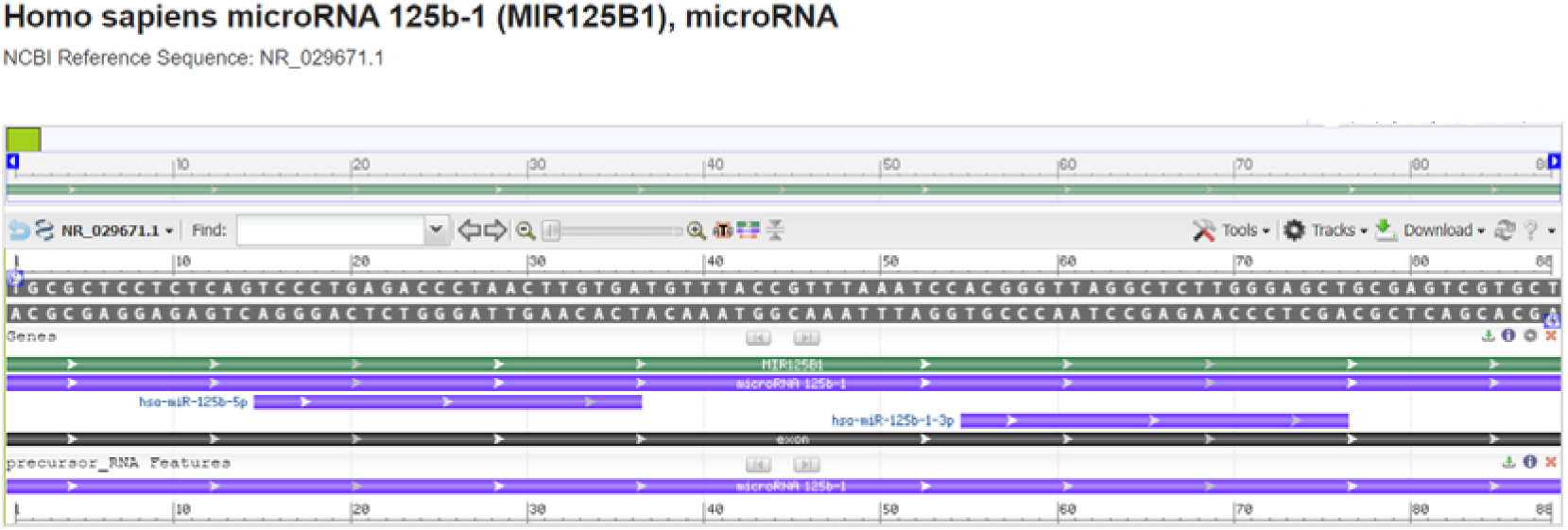
The Homo sapiens microRNA 125b-1 (MIR125B1), microRNA - *modeltemplate aligment*. **Source**: https://www.ncbi.nlm.nih.gov/nuccore/NR_029671.1?report=graph

### Molecular model of microRNA 125b-1

Based on the sequence alignment between the Homo sapiens microRNA 125b-1 nucleotide sequence: UGCGCUCCUCUCAGUCCCUGAGACCCUAACUUGUGAUGUUUACCGUUUAAAUCCACGGGUUAGGCUCUUGGGAGCUGCGAGUCGUGCU, and secondary structure: .((((..(((.((((.(((((((.(((((((((((..(((((…..))))).))))))))))).))))))).)))).)))..)))). and the template structure, the structural template for microRNA 125b-1 was produced. Assessment tools were used para measure the reliability of the designed structure. Thereby, using the RNAComposer comparative nucleotide modeling server, we generate a homology model of the Homo sapiens microRNA 125b-1 (MIR125B1), microRNA with Cartoon-style structure in pdb format (Figure 20).

**Figure 20.**
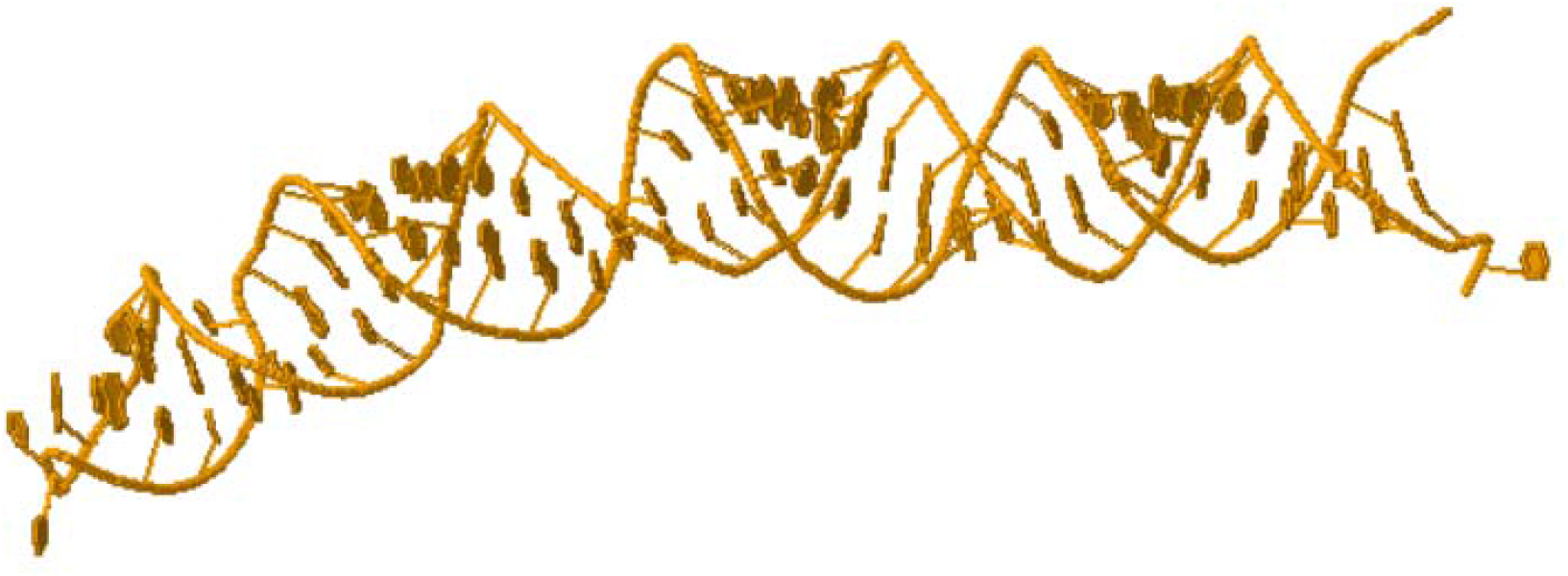
Homology model of the Homo sapiens microRNA 125b-1 (MIR125B1), microRNA **Source**: https://rnacomposer.cs.put.poznan.pl/

### Nucleotide sequence of miRNA-196b

To construct the structure of miRNA-196b the nucleotide sequences with the NCBI identifier code: NR_029911.1 were used under the FASTA format obtained from GenBank. The miRNA-196b was planned to encode a 84 bp linear ncRNA. Homo sapiens microRNA 196b (MIR196B), microRNA analysis is demonstrated in Figure 21.

**Figure 21.**
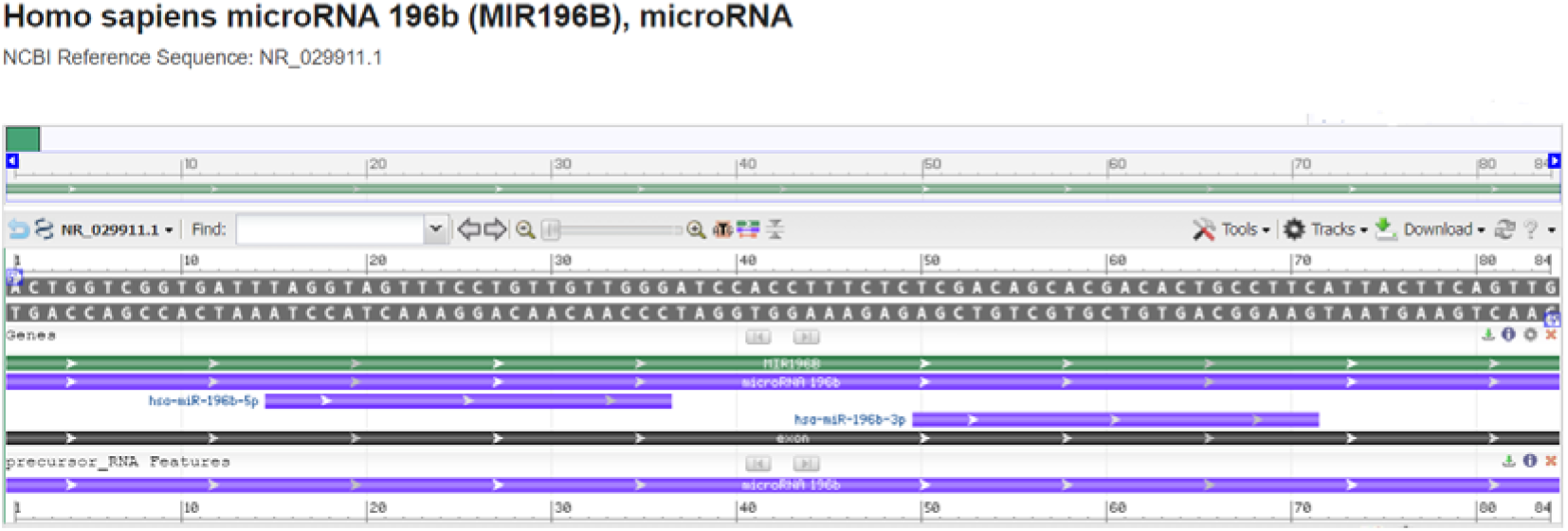
Homo sapiens microRNA 196b (MIR196B), microRNA - *model-template aligment*. **Source**: https://www.ncbi.nlm.nih.gov/nuccore/NR_029911.1?report=graph

### Molecular model of microRNA 196b

Nucleotide sequences of Homo sapiens microRNA 196b (MIR196B) were acquired employing FASTA format sequence:

ACUGGUCGGUGAUUUAGGUAGUUUCCUGUUGUUGGGAUCCACCUUUCUCUCGACAGCACGACACUGCCUUCAUUACUUCAGUUG, and secondary structure: (((((..((((((..(((((((.((.(((((((((((……….))))))))))).)).)))))))..)))))))))))..; modeling was performed employing the RNAComposer, optimized and adjusted for alignment between structural templates and microRNA 196b nucleotide. Based on sequence alignment between the template structure and microRNA nucleotide, a structural model was built for the nucleotide in question. So, employing RNAComposer of comparative nucleotide modeling we generated a homology model of Homo sapiens microRNA 196b (MIR196B), microRNA, with CPK spacefill style structure in pdb format demonstrated in Figure 22.

**Figure 22.**
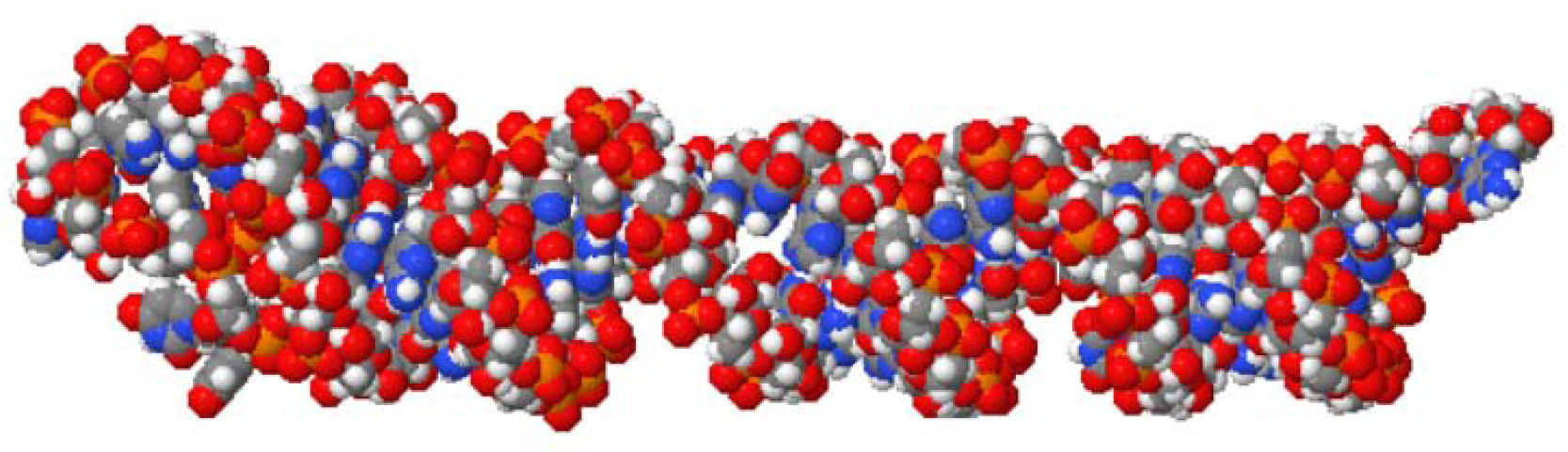
Homology model of the Homo sapiens microRNA 196b (MIR196B), microRNA **Source**: https://rnacomposer.cs.put.poznan.pl/

### Nucleotide sequence of miRNA-425

The reconstruction of miRNA-425 was performed from a nucleotide sequence archive in FASTA format obtained in GenBank database with the identifier code NCBI Reference Sequence: NR_029948.1. The miRNA-425 was planned to encode a 87 bp linear ncRNA. All coded sequences selected in FASTA format, used the annotation of the NCBI - Graphics - Homo sapiens microRNA 425 (MIR425), microRNA. The microRNA-425 analysis is shown in Figure 23.

**Figure 23.**
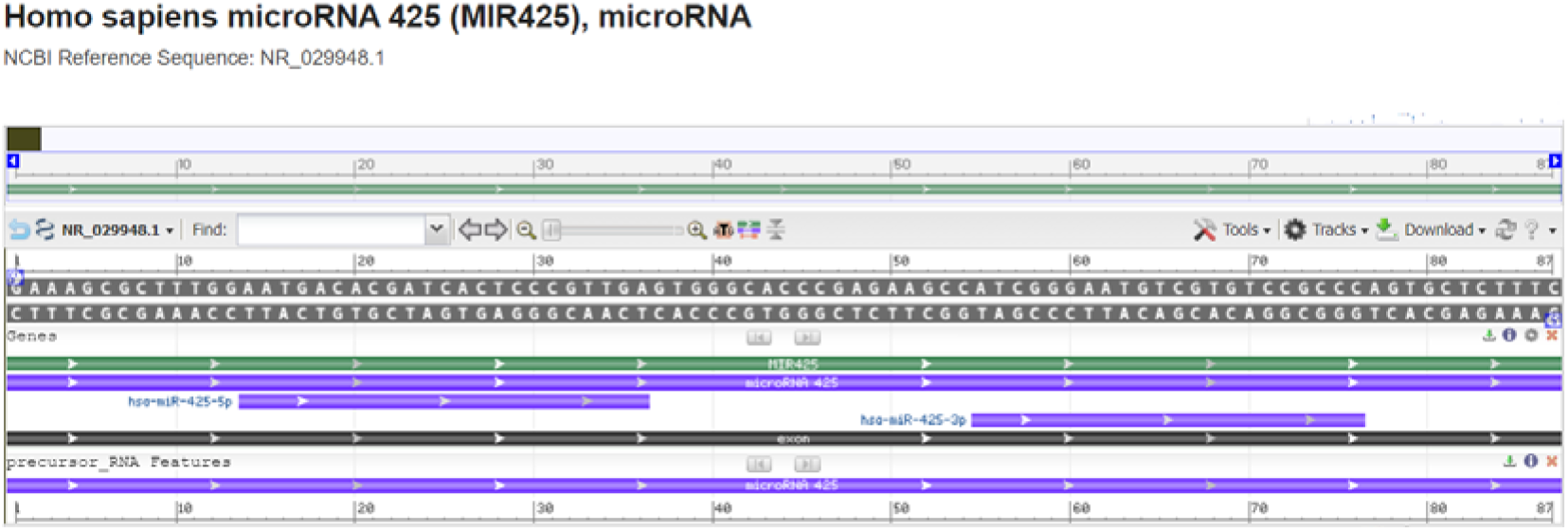
Homo sapiens microRNA 425 (MIR425), microRNA - *model-template aligment*. **Source**: https://www.ncbi.nlm.nih.gov/nuccore/NR_029948.1?report=graph

### Molecular model of microRNA-425

Based on the sequence alignment between the Homo sapiens microRNA 425 nucleotide sequence: GAAAGCGCUUUGGAAUGACACGAUCACUCCCGUUGAGUGGGCACCCGAGAAGCCAUCGGGAAUGUCGUGUCCGCCCAGUGCUCUUUC, and secondary structure: (((((.((.((((…(((((((.((.(((((…….(((………)))..))))).)))))))))…)))).)).))))), and the template structure, the structural template for microRNA-425 was produced.

Assessment tools were used para measure the reliability of the designed structure. Thereby, using the RNAComposer comparative nucleotide modeling server, we generate a homology model of the Homo sapiens microRNA 425, with Cartoon-style structure in pdb format (Figure 24).

**Figure 24.**
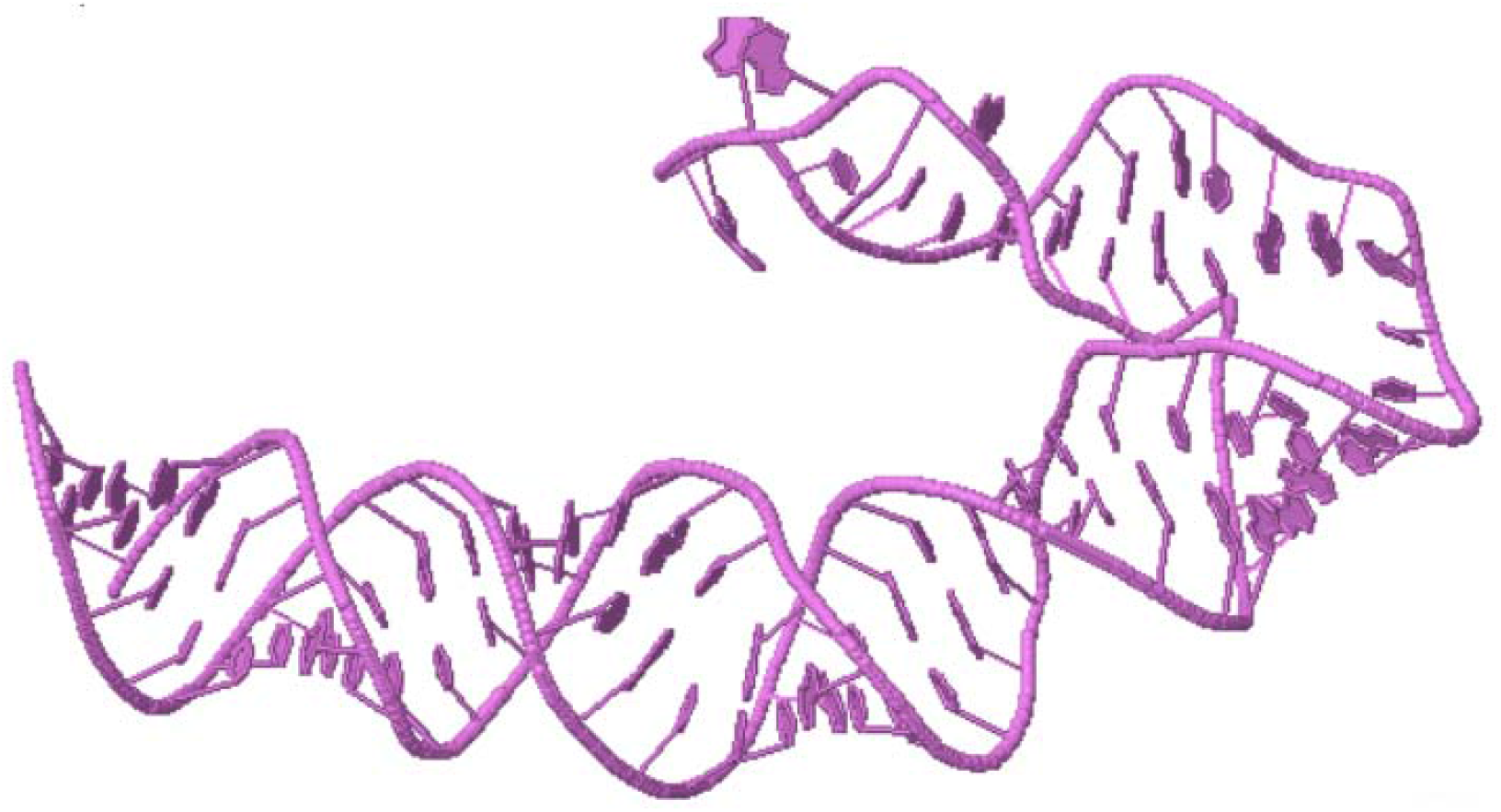
Homology model of the Homo sapiens microRNA 425 (MIR425), microRNA **Source**: https://rnacomposer.cs.put.poznan.pl/

### Nucleotide sequence of miRNA-452

To construct the structure of miRNA-452 the nucleotide sequences with the NCBI identifier code: NR_029973.1 were used under the FASTA format obtained from GenBank. The miRNA-452 was planned to encode a 85 bp linear ncRNA. Homo sapiens microRNA 452 (MIR452), microRNA analysis is demonstrated in Figure 25.

**Figure 25.**
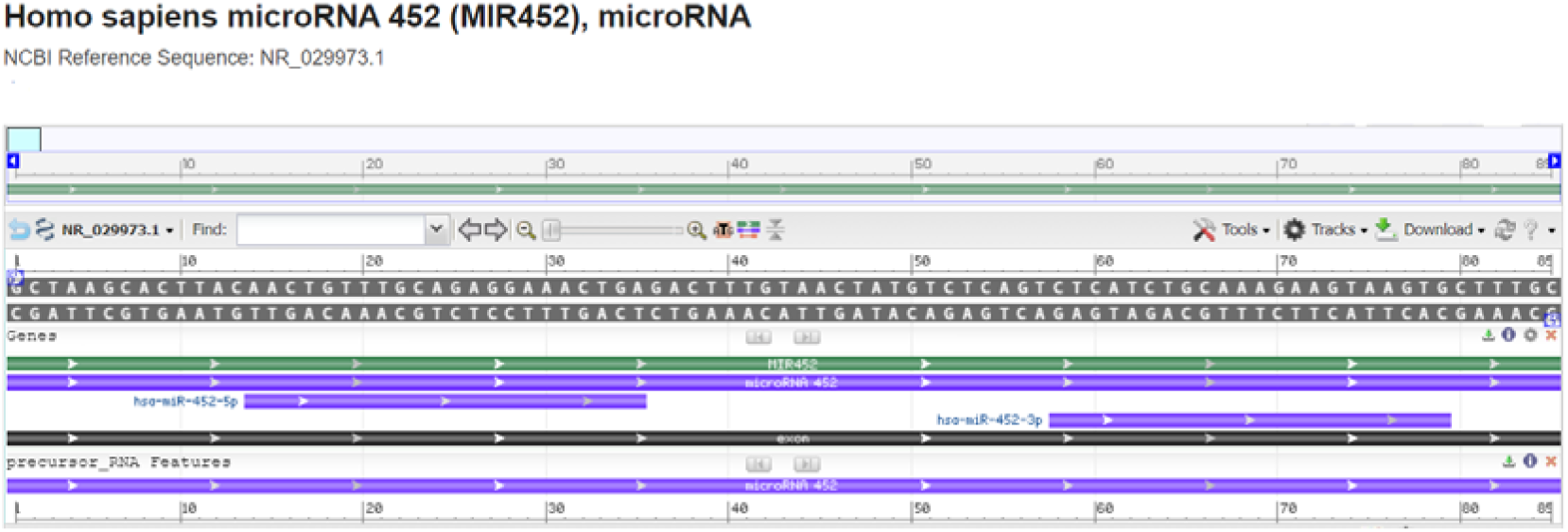
Homo sapiens microRNA 452 (MIR452), microRNA - *model-template aligment*. **Source**: https://www.ncbi.nlm.nih.gov/nuccore/NR_029973.1?report=graph

### Molecular model of miRNA-452

Nucleotide sequences of Homo sapiens microRNA 452 (MIR452) were acquired employing FASTA format sequence: GCUAAGCACUUACAACUGUUUGCAGAGGAAACUGAGACUUUGUAACUAUUCUCAGUCUCAUCUGCAAAGAAGUAAGUGCUUUGC, and secondary structure:((.((((((((((…..((((((((.((.((((((((………..)))))))).)).))))))))…)))))))))).)); modeling was performed employing the RNAComposer, optimized and adjusted for alignment between structural templates and miRNA-452 nucleotide. Based on sequence alignment between the template structure and miRNA-452 nucleotide, a structural model was built for the nucleotide in question. So, employing RNAComposer of comparative nucleotide modeling we generated a homology model of microRNA 452 (MIR452), with CPK spacefill style structure in pdb format demonstrated in Figure 26.

**Figure 26.**
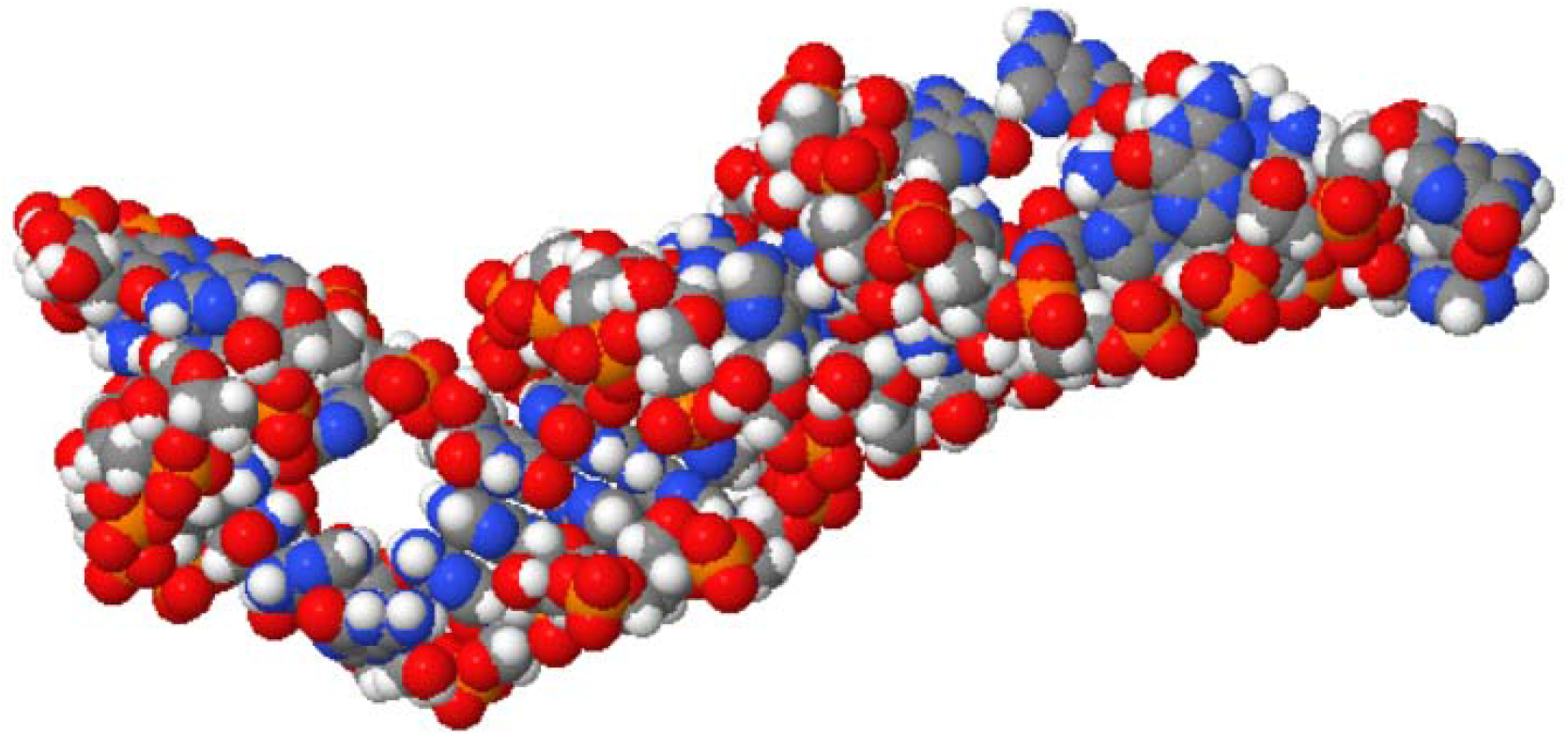
Homology model of the Homo sapiens microRNA 452 (MIR452), microRNA **Source**: https://rnacomposer.cs.put.poznan.pl/

### Nucleotide sequence of miRNA-455

The reconstruction of miRNA-455 was performed from a nucleotide sequence archive in FASTA format obtained in GenBank database with the identifier code NCBI Reference Sequence: NR_030255.1. The miRNA-455 was planned to encode a 96 bp linear ncRNA. All coded sequences selected in FASTA format, used the annotation of the NCBI - Graphics. The Homo sapiens microRNA 455 (MIR455), microRNA analysis is shown in Figure 27.

**Figure 27.**
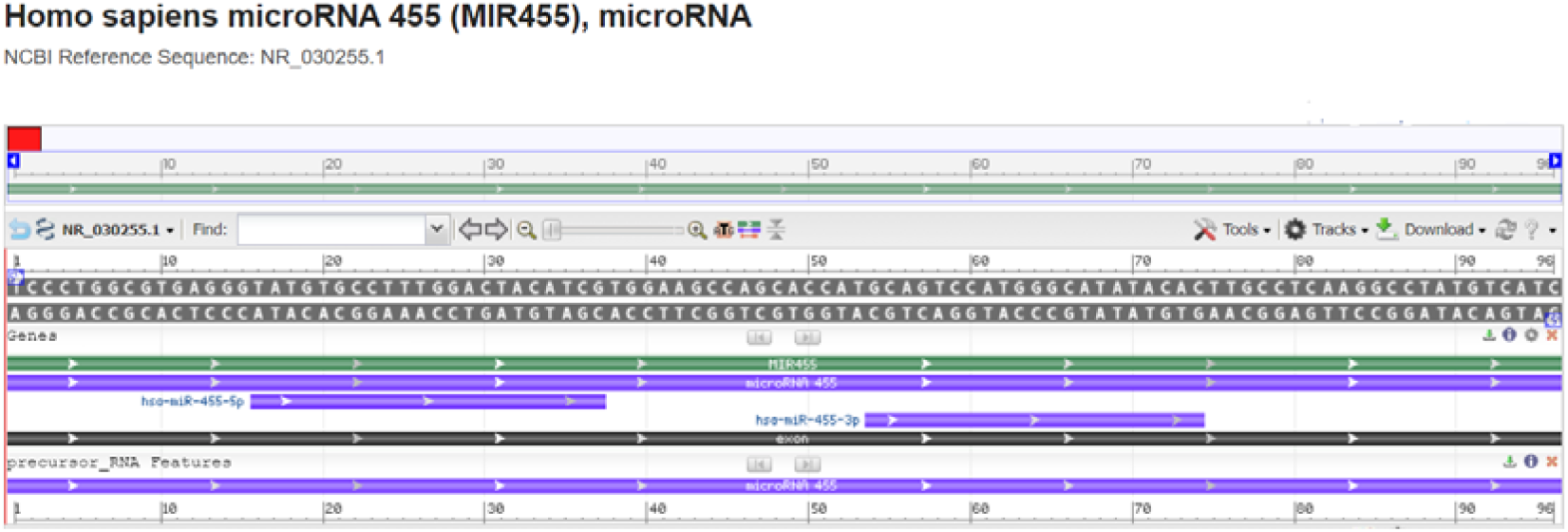
Homo sapiens microRNA 455 (MIR455), microRNA - *model-template aligment*. **Source**: https://www.ncbi.nlm.nih.gov/nuccore/NR_030255.1?report=graph

### Molecular model of microRNA-455

Based on the sequence alignment between the Homo sapiens microRNA 455 nucleotide sequence: UCCCUGGCGUGAGGGUAUGUGCCUUUGGACUACAUCGUGGAAGCCAGCACCAUGCAGUCCAUGGGCAUAUACACUUGCCUCAAGGCCUAUGUCAUC, and secondary structure: …..(((.(((((((((((((((.((((((.(((.(((……..))).))).)))))).))))))))))…..)))))..)))………, and the template structure, the structural template for microRNA 455 was produced. Assessment tools were used para measure the reliability of the designed structure. Thereby, using the RNAComposer comparative nucleotide modeling server, we generate a homology model of the Homo sapiens microRNA 455 (MIR455), microRNA with Cartoon-style structure in pdb format (Figure 28).

**Figure 28.**
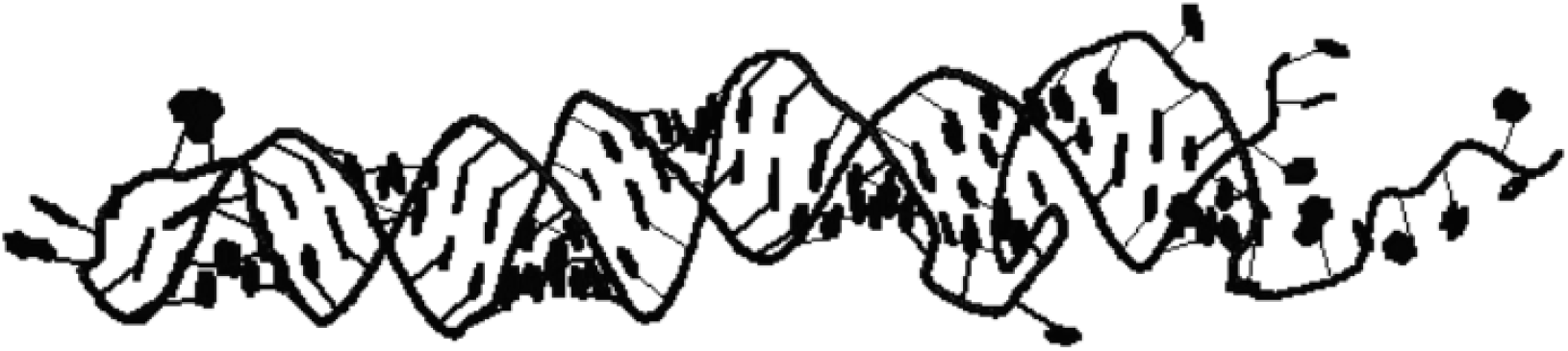
Homology model of the Homo sapiens microRNA 455 (MIR455), microRNA. **Source**: https://rnacomposer.cs.put.poznan.pl/

### Nucleotide sequence of miRNA-502

To construct the structure of miRNA-502 the nucleotide sequences with the NCBI identifier code: NR_030226.1 were used under the FASTA format obtained from GenBank. The miRNA-146B was planned to encode a 86 bp linear ncRNA. Homo sapiens microRNA 502 (MIR502), microRNA analysis is demonstrated in Figure 29.

**Figure 29.**
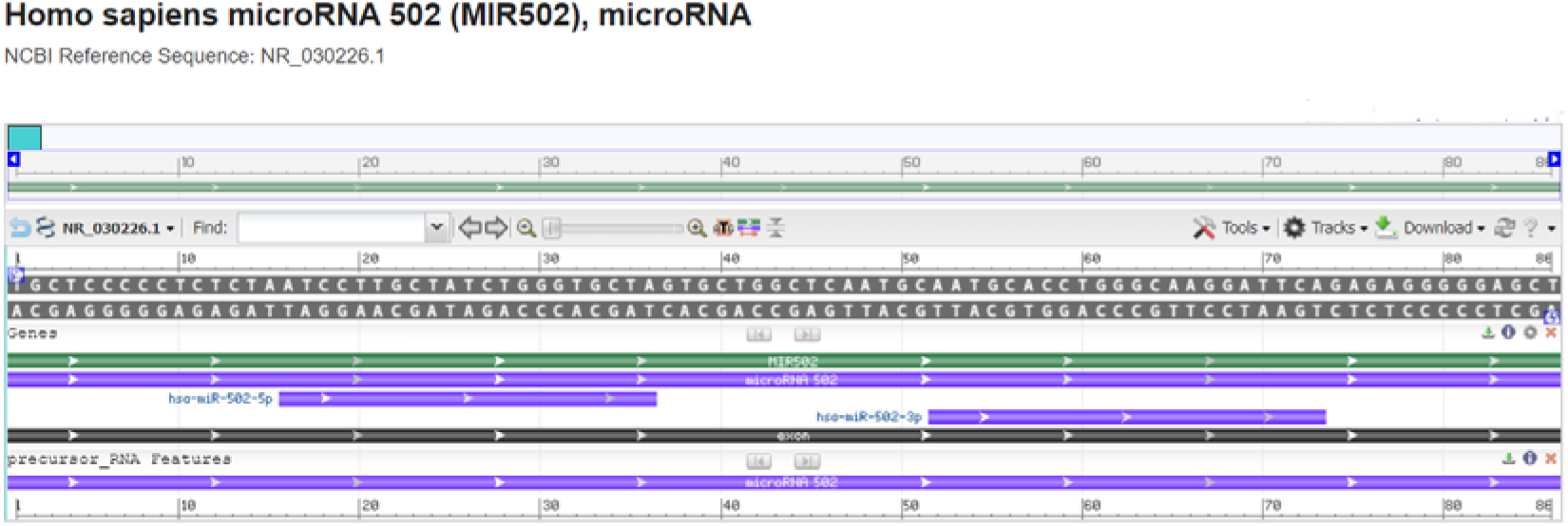
Homo sapiens microRNA 502 (MIR502), microRNA - *model-template aligment*. **Source**: https://www.ncbi.nlm.nih.gov/nuccore/NR_030226.1?report=graph

### Molecular model of microRNA-502

Nucleotide sequences of Homo sapiens microRNA 502 (MIR502), microRNA were acquired employing FASTA format sequence: UGCUCCCCCUCUCUAAUCCUUGCUAUCUGGGUGCUAGUGCUGGCUCAAUGCAAUGCACCUGGGCAAGGAUUCAGAGAGGGGGAGCU and secondary structure: .(((((((((((((((((((((((…(((((((…(((………)))..))))))))))))))))).))))))))))))).; modeling was performed employing the RNAComposer, optimized and adjusted for alignment between structural templates and miRNA-502 nucleotide. Based on sequence alignment between the template structure and miRNA-502 nucleotide, a structural model was built for the nucleotide in question. So, employing RNAComposer of comparative nucleotide modeling we generated a homology model of microRNA 502 (MIR502), with CPK spacefill style structure in pdb format demonstrated in Figure 30.

**Figure 30.**
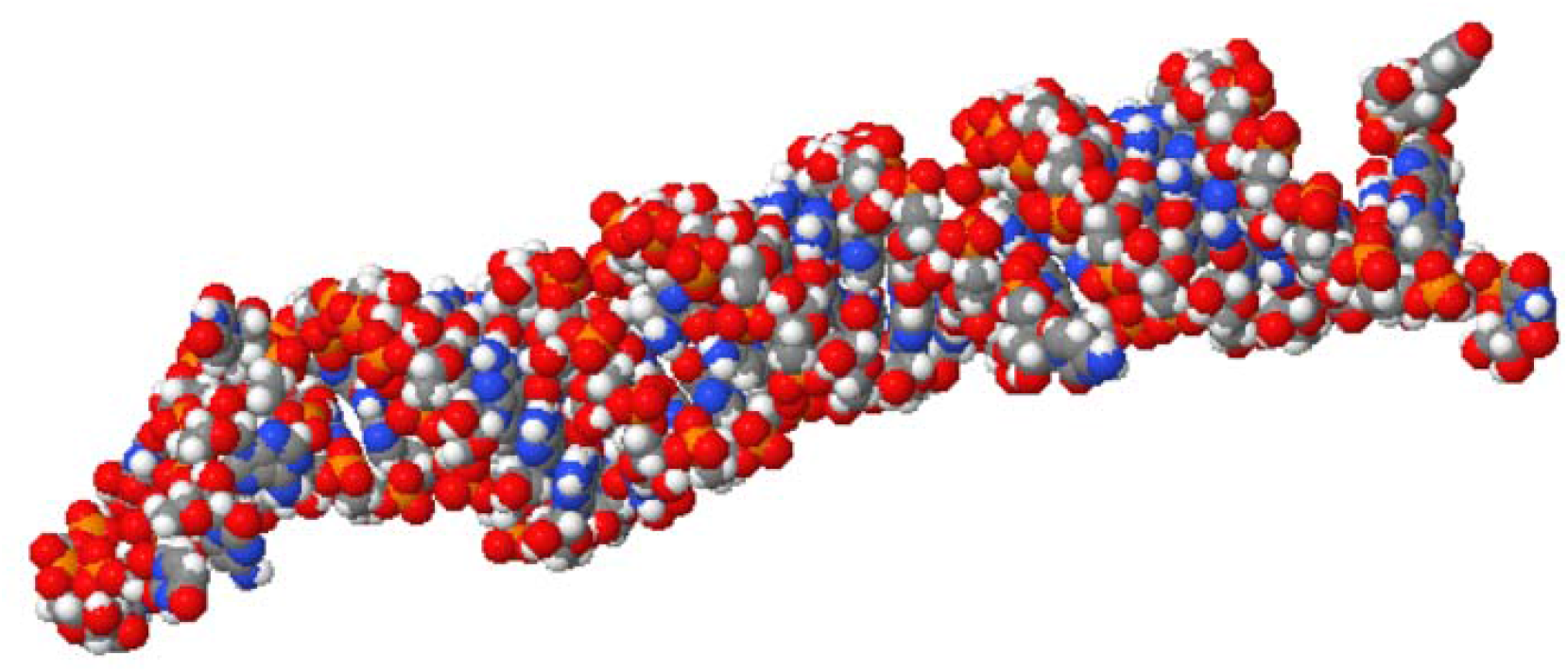
Homology model of the Homo sapiens microRNA 502 (MIR502), microRNA **Source**: https://rnacomposer.cs.put.poznan.pl/

### Nucleotide sequence of miRNA-539

The reconstruction of miRNA-539 was performed from a nucleotide sequence archive in FASTA format obtained in GenBank database with the identifier code NCBI Reference Sequence: NR_030256.1. The miRNA-539 was planned to encode a 78 bp linear ncRNA. All coded sequences selected in FASTA format, used the annotation of the NCBI - Graphics. The Homo sapiens microRNA 539 (MIR539), microRNA analysis is shown in Figure 31.

**Figure 31.**
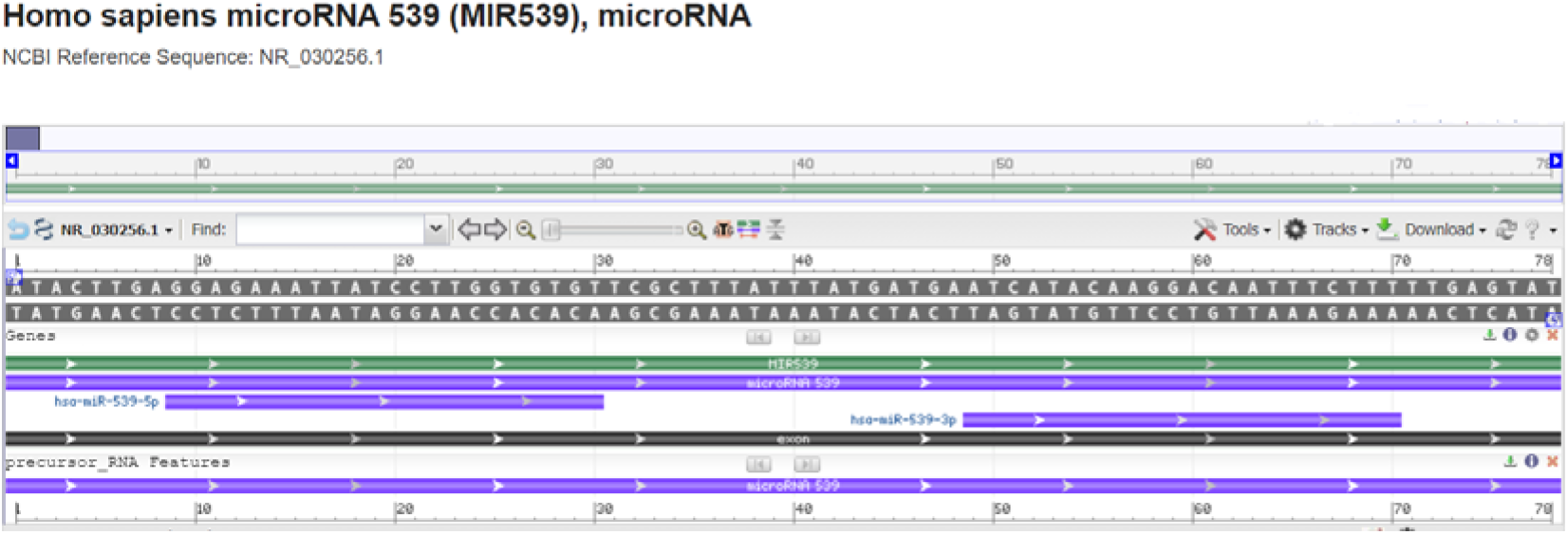
Homo sapiens microRNA 539 (MIR539), microRNA - *model-template aligment*. **Source**: https://www.ncbi.nlm.nih.gov/nuccore/NR_030256.1?report=graph

### Molecular model of microRNA-549

Based on the sequence alignment between the Homo sapiens microRNA-549 nucleotideAUACUUGAGGAGAAAUUAUCCUUGGUGUGUUCGCUUUAUUUAU GAUGAAUCAUACAAGGACAAUUUCUUUUUGAGUAU, and (((((((((((((((((.((((((.(((((((…(((….))).))).)))))))))).))))))))))))))))), and the template structure, the structural template for microRNA-549 was produced. Assessment tools were used para measure the reliability of the designed structure. Thereby, using the RNAComposer comparative nucleotide modeling server, we generate a homology model of the Homo sapiens microRNA 539 (MIR539), microRNA, microRNA with Cartoon-style structure in pdb format (Figure 32).

**Figure 32.**
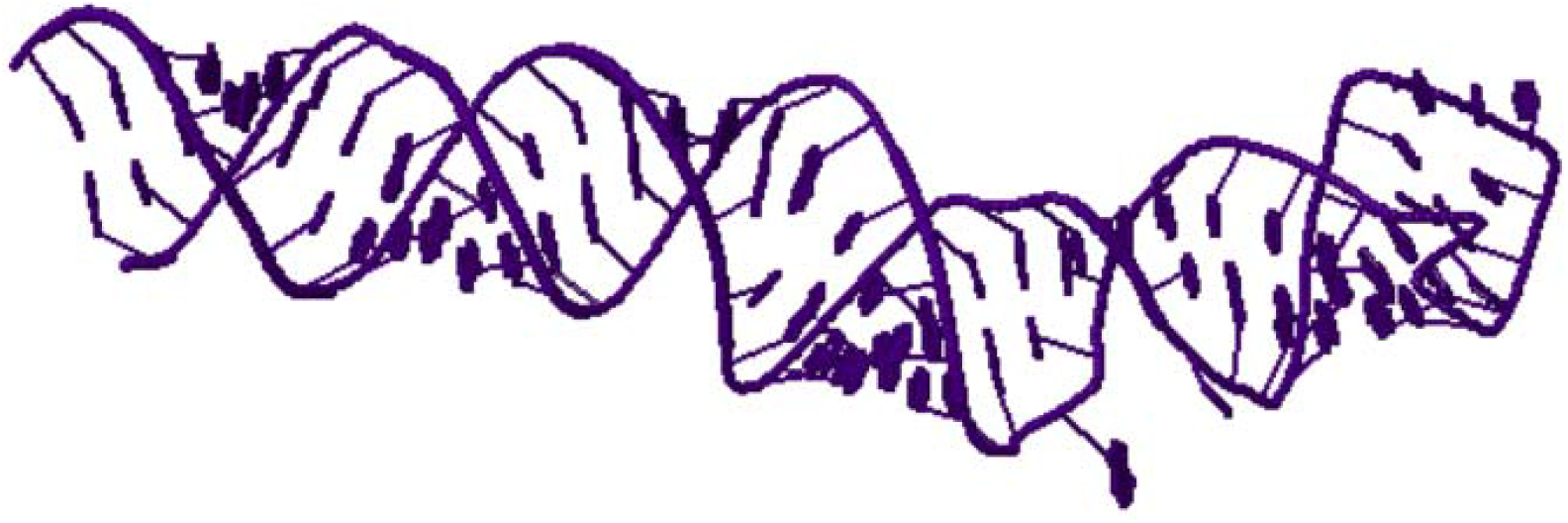
Homology model of the Homo sapiens microRNA 539 (MIR539), microRNA **Source**: https://rnacomposer.cs.put.poznan.pl/

### Nucleotide sequence of miRNA-660

To construct the structure of miRNA-660 the nucleotide sequences with the NCBI identifier code: NR_030397.1 were used under the FASTA format obtained from GenBank. The miRNA-660 was planned to encode a 97 bp linear ncRNA. Homo sapiens microRNA 660 (MIR660), microRNA analysis is demonstrated in Figure 33.

**Figure 33.**
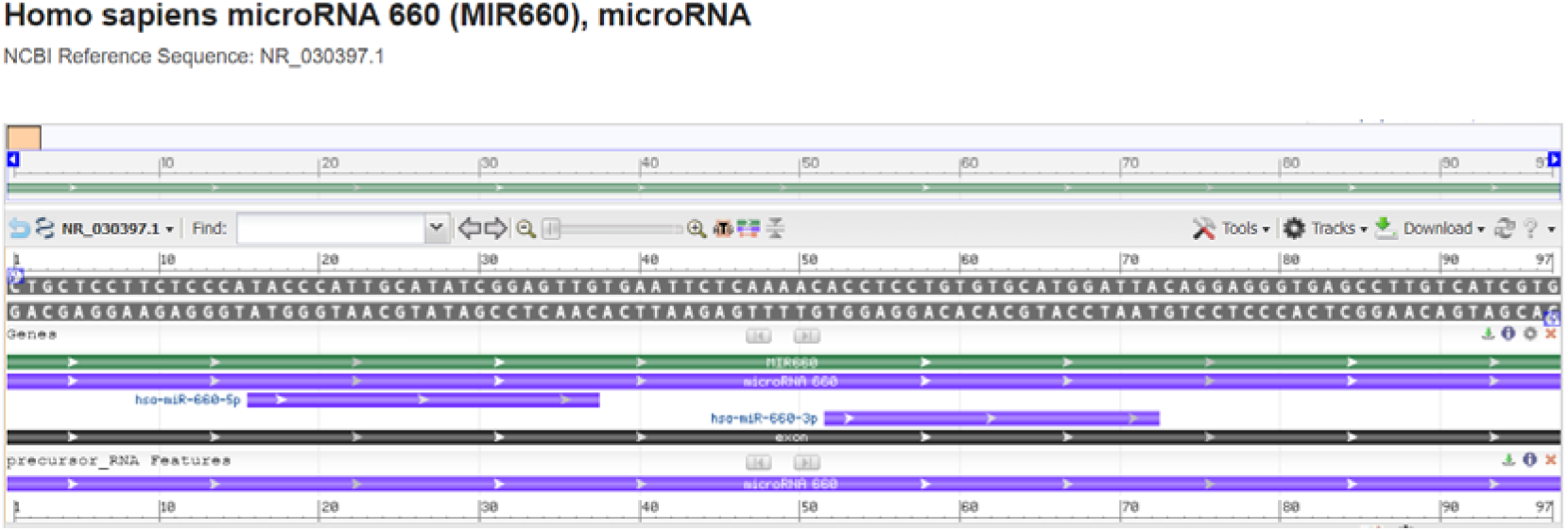
Homo sapiens microRNA 660 (MIR660), microRNA - *model-template aligment*. **Source**: https://www.ncbi.nlm.nih.gov/nuccore/NR_030397.1?report=graph

### Molecular model of miRNA-146a

Nucleotide sequences of Homo sapiens microRNA 660 (MIR660) were acquired employing FASTA format sequence CUGCUCCUUCUCCCAUACCCAUUGCAUAUCGGAGUUGUGAAUUCUCAAAACACCUCCUGUGUGCAUGGAUUACAGGAGGGUGAGCCUUGUCAUCGUG, and secondary structure ..((((.((((((…..((((.((((((.((((.(((………..))).)))).))))))))))……)))))).))))…………; modeling was performed employing the RNAComposer, optimized and adjusted for alignment between structural templates and miRNA-660 nucleotide. Based on sequence alignment between the template structure and miRNA-660 nucleotide, a structural model was built for the nucleotide in question. So, employing RNAComposer of comparative nucleotide modeling we generated a homology model of Homo sapiens microRNA 660 (MIR660), microRNA, demonstrated in Figure 34.

**Figure 34.**
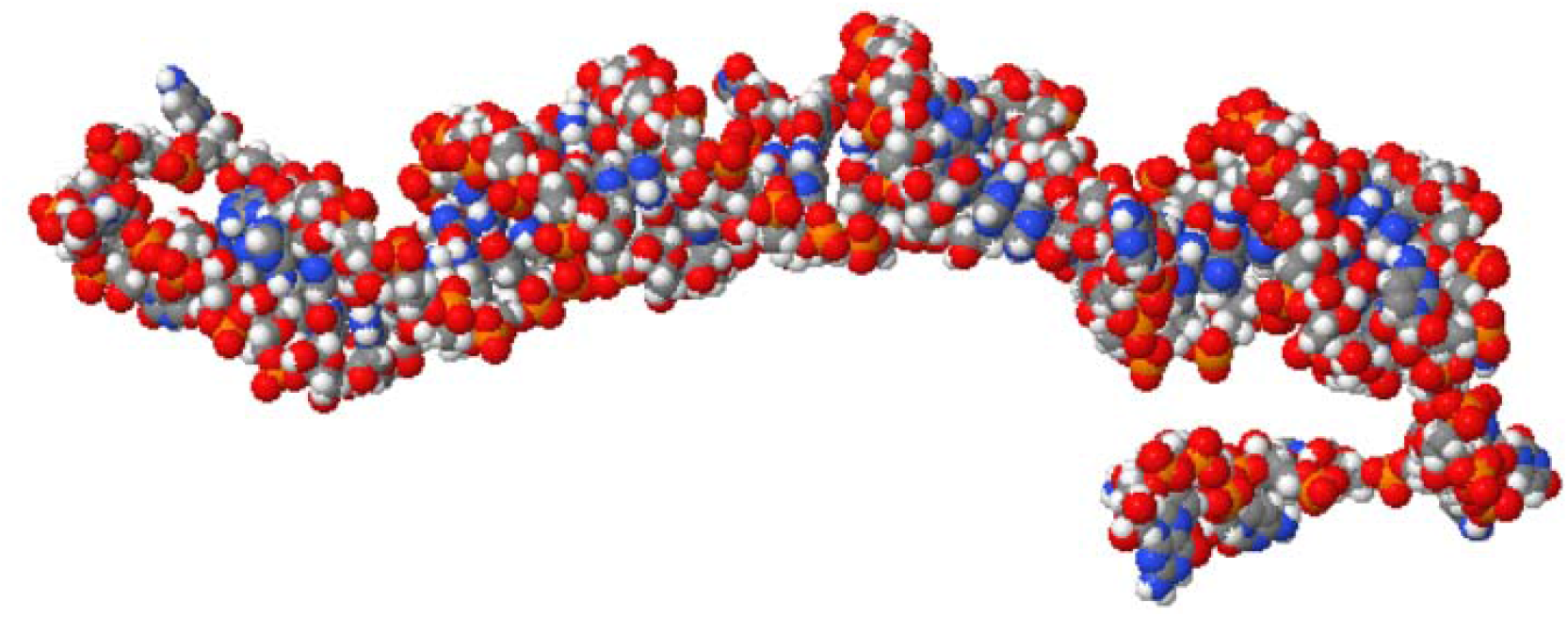
Homology model of the Homo sapiens microRNA 660 (MIR660), microRNA. **Source**: https://rnacomposer.cs.put.poznan.pl/

## DISCUSSION

miRNAs have been expressed in a number of infectious and contagious diseases, and have been proposed as potential diagnostic, prognostic and therapeutic markers in leprosy. It has been shown that M. leprae, controls in dysregulation of miRNAs at the site of the infectious lesion in leprosy patients and participating in the host therapeutic response.^6^

The miRNAs are short noncoding RNAs that intervene in post-transcription command of gene manifestation in multicellular beings, controlling role in multiples physiological and pathophysiological functions. Studies and the comprehension about the function and structure of miRNAs have grown significantly in recent years. Our study presents a tutorial on molecular modeling and demonstrates *in silico* the projection of the molecular structure of 17 miRNAs expressed in leprosy, drawing *in silico* their molecular structures.

Leprosy is a chronic infectious disease which has as etiologic agent M. leprae, being a public health problem in the American, African and Asian continents, especially in developing countries and especially in poor countries.^7^ M. leprae is an intracellular bacillus which has a long incubation time, slow proliferation, and peripheral nerve tropism. Leprosy has three distinct forms, tuberculoid, lepromatous and an intermediate form.^8^

Various microRNAs and their target genes participate of tissue damage in leprosy, and based on literature review studies, we address the following miRNAs most frequently expressed in the leprosy lesion: miRNA-26a, miRNA-27a, miRNA-27b, miRNA-29c, miRNA-34c, miRNA-92a-1, miRNA-99a-2, miRNA-101-1, miRNA-101-2, miRNA-125b, miRNA-196b, miRNA-425, miRNA-452, miRNA-455, miRNA-502, miRNA-539, and miRNA-660.

The combination of several miRNAs as biomarkers is considered as a more efficient option due to the possible overlap in miRNA targeting.^9^ The combination of 17 miRNAs evaluated in a study comparing normal individuals with individuals with leprosy showed 80% sensitivity and 91% specificity in discriminating between leprosy and healthy individuals indicating a high diagnostic potency of the miRNAs described above.^10^

With the advancement of bioinformatics several tools have been developed for each step of miRNA biogenesis, helping researchers in molecular biology investigations. Bioinformatics tools for miRNAs design have been showing accessibility for experimental evaluations and structure definitions. The computational study of miRNAs is fundamentally based on the performance analysis of their primary and secondary structures. Reviewing the available literature, we found no studies with the *in silico* construction of miRNA-26a, miRNA-27a, miRNA-27b, miRNA-29c, miRNA-34c, miRNA-92a-1, miRNA-99a-2, miRNA-101-1, miRNA-101-2, miRNA-125b, miRNA-196b, miRNA-425, miRNA-452, miRNA-455, miRNA-502, miRNA-539, and miRNA-660 with a 3-D structure model.

An alternative method using three dimensions (3D) graphical presentation of secondary structures of miRNAs is a promising method as it creates data matrices based on structural information. In our study, the sequence analysis of selected miRNAs was evaluated and 3D structural models were designed. The nucleotide analysis of miRNAs was performed in FASTA format, and the 3D structuring was performed using RNAComposer. Thus, to construct the 3D structural models of the miRNAs, a strategy of structural analysis by homology with the sequence from the Nucleotide database was used. The use of molecular models can promote progress in the development of new drugs that facilitate therapy in target organs.

## CONCLUSION

We demonstrate *in silico* design of selected molecular structures of 17 miRNAs expressed in leprosy through computational biology.

The function and structure of miRNAs are determined by nucleotide sequences and methods for the early knowledge of these structures can allow the identification and the localization of binding sites on nucleotides which can be of fundamental for clinical and therapeutic use.

